# Host-derived CEACAM-laden vesicles engage enterotoxigenic *E. coli* for elimination and toxin neutralization

**DOI:** 10.1101/2024.07.24.604983

**Authors:** Alaullah Sheikh, Debayan Ganguli, Tim J. Vickers, Bernhard Singer, Jennifer Foulke-Abel, Marjahan Akhtar, Nazia Khatoon, Bipul Setu, Supratim Basu, Clayton Harro, Nicole Maier, Wandy L. Beatty, Subhra Chakraborty, Tafiqur R. Bhuiyan, Firdausi Qadri, Mark Donowitz, James M. Fleckenstein

## Abstract

Enterotoxigenic *Escherichia coli* (ETEC) cause hundreds of millions of diarrheal illnesses annually ranging from mildly symptomatic cases to severe, life-threatening cholera-like diarrhea. Although ETEC are associated with long-term sequelae including malnutrition, the acute diarrheal illness is largely self-limited. Recent studies indicate that in addition to causing diarrhea, the ETEC heat-labile toxin (LT) modulates the expression of many genes in intestinal epithelia, including carcinoembryonic cell adhesion molecules (CEACAMs) which ETEC exploit as receptors, enabling toxin delivery. Here however, we demonstrate that LT also enhances the expression of CEACAMs on extracellular vesicles (EV) shed by intestinal epithelia and that CEACAM-laden EV increase in abundance during human infections, mitigate pathogen-host interactions, scavenge free ETEC toxins, and accelerate ETEC clearance from the gastrointestinal tract. Collectively, these findings indicate that CEACAMs play a multifaceted role in ETEC pathogen-host interactions, transiently favoring the pathogen, but ultimately contributing to innate responses that extinguish these common infections.

**Significance statement:** Enterotoxigenic *E. coli,* characterized by the production of heat-labile (LT) and heat-stable (ST) toxins, are a very common cause of diarrhea in low-income regions responsible for hundreds of millions of infections each year, and the major cause of diarrhea in travelers to endemic areas. Although these infections may be severe and cholera-like, they are typically self-limited. These studies demonstrate that extracellular vesicles produced by host intestinal cells can capture the bacteria and its secreted toxins at a distance from the cell surface, potentially acting as molecular decoys to neutralize the enterotoxins and extinguish the infection.

## Introduction

Enterotoxigenic *Escherichia coli* (ETEC) comprise a diverse diarrheagenic pathovar defined by the production of heat-labile (LT) and/or heat-stable (ST) enterotoxins. These pathogens are thought to account for hundreds of millions of cases of diarrheal illness annually with young children in low-middle income countries disproportionately affected^1^. ETEC have remained a leading cause of death due to acute diarrheal illness^2^, and are associated with long-term sequelae including malnutrition, growth stunting^3,4^ and cognitive impairment^5^.

The basic mechanism by which these pathogens cause diarrheal illness is well-established. Heat-labile toxin (LT) binds to gangliosides on the intestinal surface, and once internalized stimulates production of cAMP. Heat-stable toxins (ST) bind to guanylate cyclase C on the surface of enterocytes to stimulate production of cGMP. These cyclic nucleotides, cAMP and cGMP, in turn activate protein kinase A (PKA) and protein kinase G (PKG), respectively. Kinase -mediated phosphorylation of cellular ion channels including the cystic fibrosis transmembrane regulator (CFTR), and the sodium hydrogen exchanger (NHE3) modulates ion transport resulting in the net export of NaCl and water into the intestinal lumen leading to watery diarrhea^6^.

Diarrheal illness caused by ETEC can range from mild to severe and cholera-like. Indeed, ETEC were initially discovered in patients with *Vibrio cholerae*-negative clinical cholera, and severe ETEC is clinically indistinguishable from cholera^7–11^. Importantly, while ETEC infections can occasionally cause more protracted symptoms^12^, acute diarrhea caused by these pathogens is typically self-limited, with resolution after several days. However, what dictates the self-limited nature of ETEC diarrhea is unknown.

To cause diarrhea, ETEC must transit to the small intestine, migrate through intestinal mucin^13^, and directly engage the brush border of enterocytes to effectively deliver toxin directly at the epithelial surface^14^. ETEC employ both plasmid-encoded adhesins unique to the ETEC pathovar^15^ as well as highly conserved chromosomally-encoded type 1 fimbriae to engage enterocytes^16^.

Recent studies demonstrate that ETEC use type 1 fimbriae^16^ to bind to members of a family of extracellular glycoproteins known as carcinoembryonic antigen related cell adhesion molecules (CEACAMs) on the surface of enterocytes, and that these interactions play a critical role in bacterial adhesion and toxin delivery to small intestinal epithelia^17^. Moreover, we found that heat-labile toxin accelerates production of CEACAMs by small intestinal enterocytes^17^, effectively modifying the epithelial landscape to transiently benefit the pathogen.

Notably however, CEACAMs are normally present in abundance in human stool^18,19^, with approximately 50-70 mg of carcinoembryonic antigen shed in the course of a day^19^. The majority of fecal CEACAMs are membrane-bound^19^, and can be released in soluble form with phosphatidylinositol specific phospholipase C (PI-PLC).

Our current studies suggest that CEACAMs play opposing roles in ETEC interactions with gastrointestinal epithelia. While initial expression of these molecules on enterocytes facilitates ETEC-host cell engagement and toxin delivery^17^, we demonstrate here that the host may interrupt these encounters by deploying CEACAM-laden extracellular vesicles as decoys to mitigate effective attachment of the bacteria to the epithelial surface while absorbing and neutralizing secreted ETEC enterotoxins, potentially explaining the self-limited nature of these common infections.

## Results

### CEACAM expression alters kinetics of ETEC intestinal colonization

Carcinoembryonic cell adhesion molecules (CEACAMs) are host cell glycoproteins comprising a large subgroup of the immunoglobulin superfamily that form homodimeric intercellular adhesion complexes^20^ and which participate in intracellular signaling pathways that can direct cellular differentiation^21^. CEACAMs differ significantly between mice and humans. Although mice have at least 20 putative CEACAM genes, only CEACAM1, CEACAM16, CEACAM18, CEACAM19, and CEACAM20 are shared with humans^22^, and all of the GPI-anchored gastrointestinal CEACAM molecules, CEACAMs 5,6, and 7, as well as CEACAM3 which is predominantly on neutrophils, are absent in conventional mice.

Therefore, we challenged CEABAC10 transgenic mice^23^, which express human CEACAMs 5-7 in the intestine, as well as CEACAM3 to examine the impact on ETEC-host interactions. These studies demonstrated that while gastrointestinal CEACAM expression was associated with modest increases in intestinal colonization immediately after ETEC (jf876, supplemental table 1) challenge (figure 1A, supplemental figure 1A), CEABAC10 mice consistently cleared ETEC more rapidly (figure 1B) than parental C57BL/6NCrl controls suggesting that these molecules play a complex role in directing the kinetics of intestinal colonization.

**figure 1.**
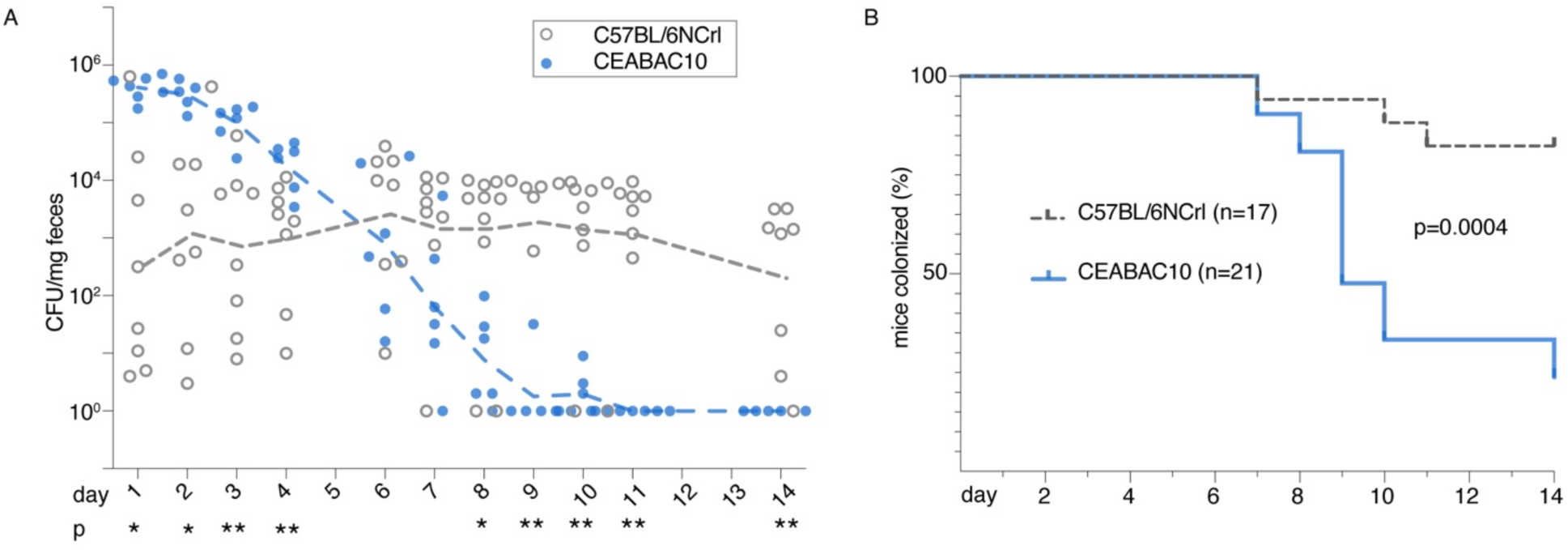
CEACAM expression alters kinetics of ETEC intestinal colonization. **A.** ETEC shed in stool following challenge of either C57BL/6NCrl control mice (n=8) or CEABAC10 CEACAM-expressing mice (n=6). Dashed lines connect geometric means. *<0.05, **<0.01 Mann Whitney two-tailed nonparametric comparison between groups. **B.** Proportion of mice remaining colonized (≥ 1 CFU/mg stool) based on fecal shedding data. Shown are combined results of two independent experiments with total of n=17 control (C57BL/6NCrl) and n=21 CEABAC10 mice. p=0.0004 Log-rank (Mantel-Cox) comparison of survival curves.

Examination of intestinal tissues of CEABAC10 transgenic mice revealed that while CEACAMs were expressed on small intestinal mucosal surfaces (figure 2A), we also found many clusters of CEACAM-positive material in the intestinal lumen, (figure 2B-C), Many of the smaller CEACAM-positive structures in the intestinal lumen were in direct contact with the bacteria (inset, figure 2C), suggesting that they may engage ETEC at a distance from the mucosal surface, potentially preventing direct interaction with epithelia. Although CEABAC10 mice may express CEACAM3 on neutrophils^23^, we were unable to demonstrate neutrophilic infiltration beyond the basolateral surface of the intestines of infected mice (supplemental figure 1B). Many of the smaller structures ∼ 100-300 nm in diameter in the lumen were similar in size to plasma-membrane derived extracellular vesicles (EV)^24,25^. Indeed, we were able to identify multiple individual CEACAM-laden EV (figure 2D), as well as clusters of vesicles (figure 2E), and direct interaction of these EV with bacteria in samples from the ileal lumen of H10407-challenged CEABAC10 mice (figure 2F). Examination of fecal material likewise revealed abundant CEACAM+ EV (figure 2G). Following challenge of CEACAM-expressing mice with ETEC expressing green fluorescent protein (GFP) (<u>jf2450,</u> supplemental table 1), we observed that while individual bacteria shed in stool appeared to be positive for CEACAMs (figure 2H), and we identified CEACAM+ EV bound to ETEC by immunogold transmission electron microscopy of fecal material following challenge (supplemental figure 2), the majority of ETEC emerged in large clusters of bacteria embedded in a CEACAM matrix (figure 2I), (<u>supplemental movie 1</u>).

**figure 2.**
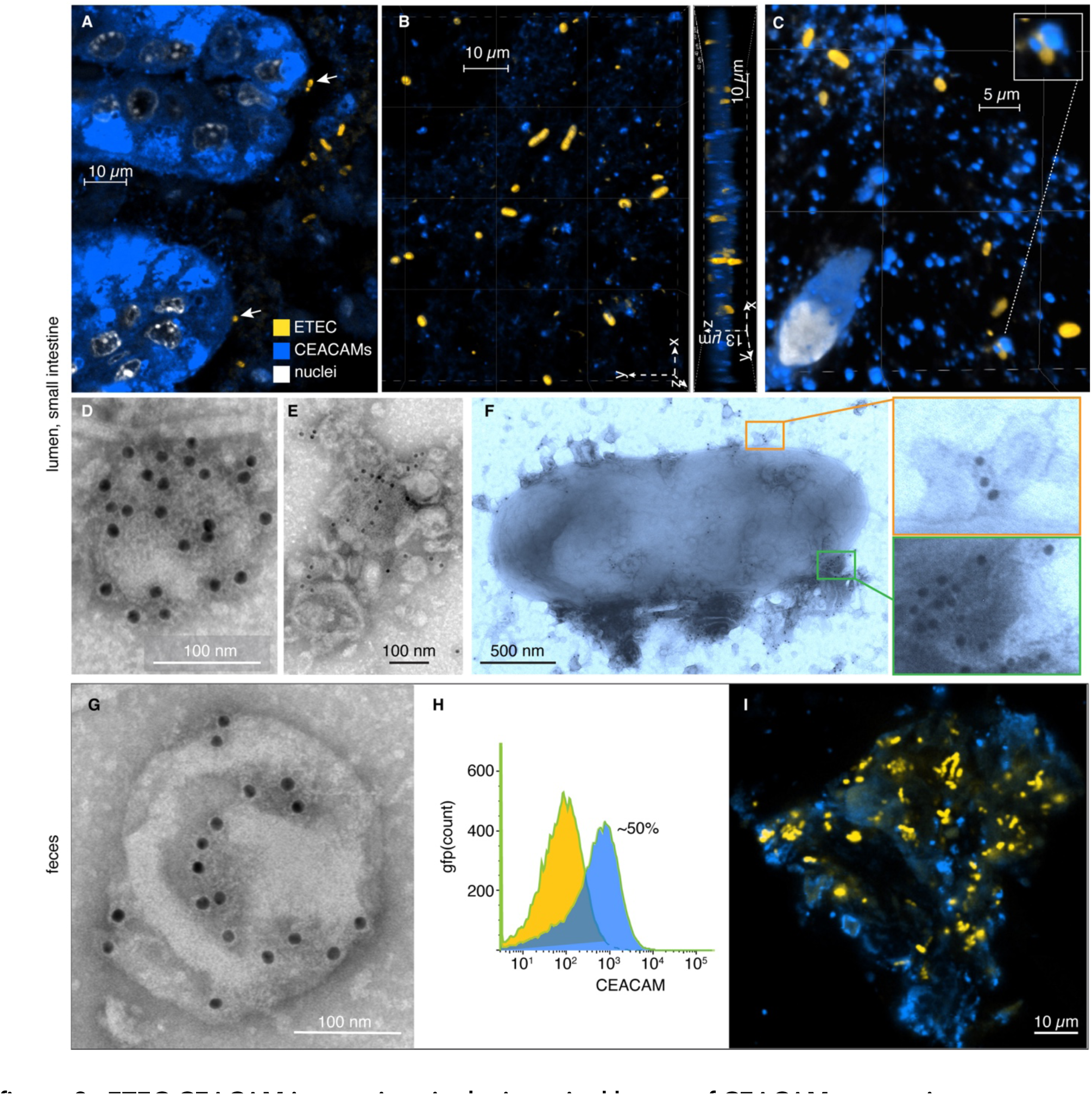
ETEC-CEACAM interactions in the intestinal lumen of CEACAM-expressing transgenic mice. Shown are confocal microscopy images of (**A**) ETEC H10407 attached to small intestinal villus enterocytes (arrows) and to CEACAM+ material in the lumen. (**B**). ETEC in the lumen reside in a CEACAM+ matrix. (**C**) Small (∼100-300 nm) CEACAM+ structures engage ETEC in the intestinal lumen. Inset shows ETEC surrounded by CEACAM + structures in the lumen. In each panel anti-CEA antibodies were used to identify CEACAMs (blue) and anti-O78 antibodies were used to identify ETEC H10407 (yellow, serotype O78). **D,E**. CEACAM6-positive vesicle, and clusters of EV isolated from ileum of CEABAC10 mouse. **F**. CEACAM6+ EV clustered on the surface of bacteria isolated from ileal lavage following H10407 challenge. **G**. CEACAM6-positive EV isolated from CEABAC10 mouse feces shown by immunogold labeling of anti-CEACAM monoclonal (9A6). **H.** Flow cytometry of GFP+ bacteria isolated from fecal resuspension supernatants following challenge of CEABAC10 mice with H10407(pGFPmut3.1), showing the proportion of GFP+ bacteria that co-labeled with CEACAMs (blue) vs those which remain unlabeled with CEACAMs (yellow). **I.** Majority of ETEC shed in feces are eliminated in large clusters of CEACAMs. Panel represents individual Z-stack confocal image of fecal resuspension. (GFP+ bacteria pseudo-colored yellow).

### CEACAMs serve as ETEC decoys and enterotoxin scavengers

Interestingly, while our earlier studies demonstrated that CEACAM6 could be identified on the microvillus surface of enterocytes where it served as a receptor for ETEC^17^, transmission electron micrographs of ETEC-infected ileal enteroid monolayers demonstrated large clusters of CEACAM-positive extracellular vesicles (EVs) interposed between the bacteria and microvilli of the intestinal brush border (figure 3A<u>,B</u>). CEACAM6-positive concentrated culture supernatants from polarized human ileal monolayers significantly blocked ETEC adhesion to target intestinal cells, while subtractive absorption with anti-CEACAM antibody partially restored effective adhesion of wild type ETEC to target intestinal epithelial cells further suggesting that CEACAMs can modulate ETEC-host interactions (figure 3C). To determine whether EV could specifically interrupt effective interaction of ETEC with target receptors on enterocytes, we first purified EVs from supernatants of small intestinal enteroid monolayers by size exclusion chromatography (supplemental figure 3a-b), and confirmed the presence of CEACAMs by immunogold labeling (supplemental figure 3c). These CEACAM+ purified vesicles adhered to the surface of ETEC (supplemental figure 4A). Interestingly, while earlier studies suggested that EV, from rat small intestine, impaired the growth of both commensal *E. coli* as well as another *E. coli* pathovar (EPEC) ^26^, EV isolated from polarized human small intestinal epithelial enteroids had no apparent impact on ETEC growth or survival (supplemental figure 4B<u>,C</u>). However, exogenous administration of these EVs significantly impaired the ability of ETEC to bind to intestinal epithelial cells (figure 3d). We had previously shown that LT increases CEACAM6 expression on the surface of intestinal enterocytes^17^. Notably, CEACAM6 abundance in EV also increased following exposure to LT and we found that EV obtained from LT-treated ileal monolayers were significantly more effective in blocking ETEC interaction with epithelial cells (figure 3E). Similarly, EV isolated from CEACAM-expressing mice were more effective in preventing bacterial adhesion compared to those from control mice (figure 3F).

**Figure 3.**
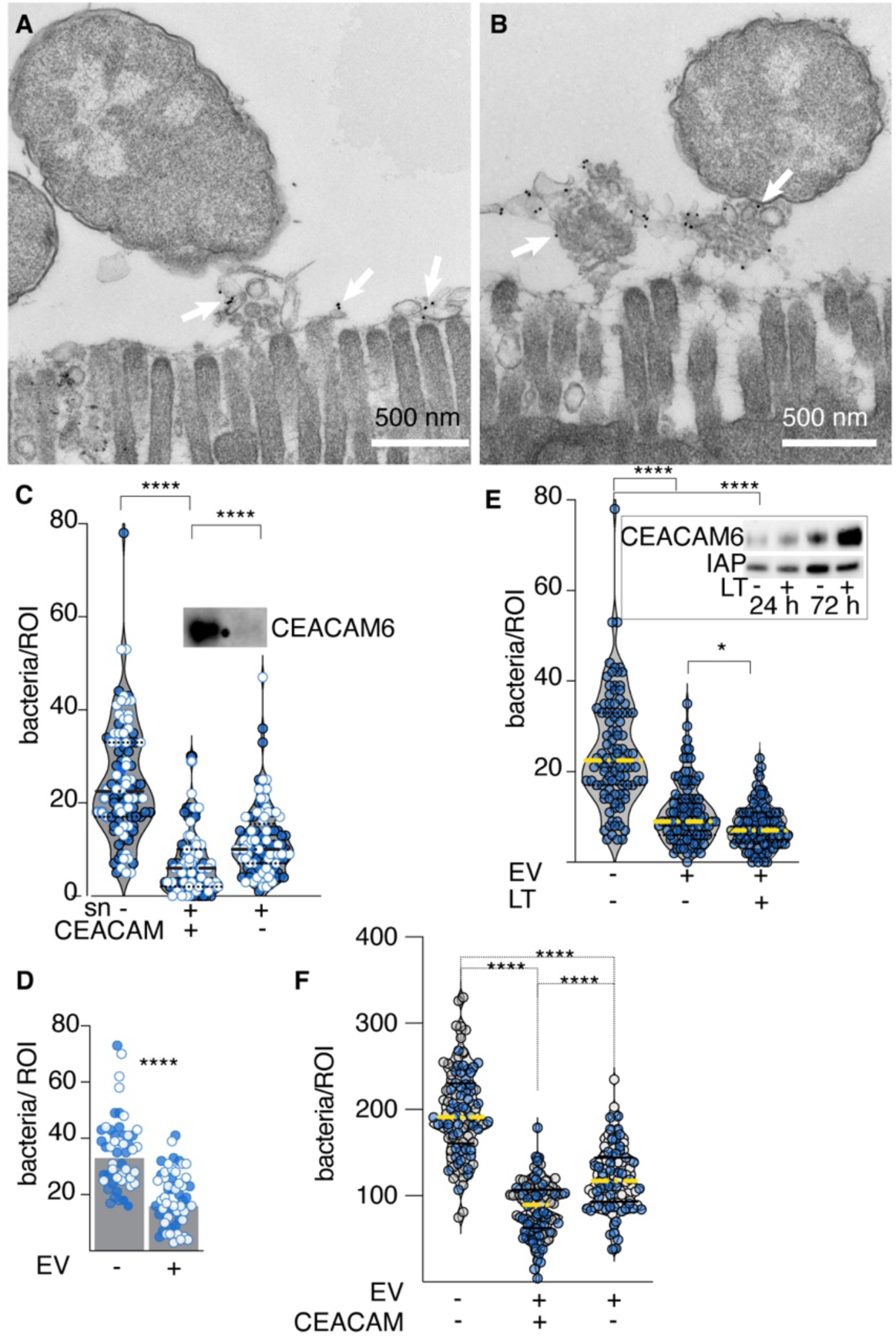
CEACAM-laden EVs block ETEC enterocyte engagement. **A**. clusters of CEACAM+ extracellular vesicles (EVs) are interposed between the brush border of small intestinal enterocytes and ETEC. **B**. CEACAM+ EVs emerging at the surface of microvilli engage ETEC. (Arrows in A,B indicate immunogold labeling of CEACAM6) **C.** Concentrated supernatants (sn) from polarized small intestinal enteroid monolayers impair ETEC pathogen-host interaction. Columns at left, middle and right of graph indicate no treatment (sn-), treatment with concentrated supernatant (sn+, CEACAM+), and after supernatant absorption of CEACAMs (sn+, CEACAM-). Shown are the results of two independent experiments. ****=p<0.0001 (Kruskall-Wallis). Horizontal lines indicate median and quartiles. Inset immunoblot indicates CEACAM6 before and after absorption. **D.** EVs purified from small intestinal enteroids inhibit attachment of ETEC to target small intestinal epithelia. Shown are the results of 2 replicate experiments. “-“= untreated ileal enteroids (n=50, total); Data reflect bacteria per region of interest (ROI) for wells without (-) and with (+) exogenously added purified EV (n=55 total). ****<0.0001 Mann-Whitney two-tailed nonparametric comparisons. **E**. LT treatment of enteroids enhances CEACAM production on EVs and EV-mediated inhibition of ETEC adhesion to target Caco-2 cells **** <0.0001, *0.02 (Kruskal-Wallis). Inset immunoblot shows impact of LT treatment on CEACAM6 expression on EV isolated after treatment for 24, 72 hours. Intestinal alkaline phosphatase (IAP) is shown as a loading control. **F**. EV isolated from murine intestine inhibit ETEC adhesion. Shown are confocal imaging data of ETEC adherent to Caco-2 cells in the absence of EV, EV isolated from CEABAC10 mice (CEACAM+), and parental C57BL6 mice (CEACAM-). ****<0.0001 (Kruskal-Wallis). Total of n=100 fields from 2 independent experiments.

We also found that purified EV can bind to heat-labile toxin (figure 4A-C, supplemental figure 5A), and were able to block LT-mediated activation of cAMP in target intestinal cells (figure 4D) suggesting that EV also bear GM1 ganglioside receptors for the LT-B subunit. Indeed, EV effectively competed with target epithelial cells resulting in complete abrogation of toxin delivery to target intestinal epithelia by wild type ETEC (figure 4E). Notably, analysis of CEACAM6+ EV fractions from size exclusion chromatography (SEC) revealed that these EV also bound fluorescently labeled cholera toxin B subunit (CT-B) (supplemental figure 5A), which like LT binds to GM-1 gangliosides. ETEC outer membrane vesicles (OMV) are known to have significant amounts of LT^27,28^ which can deliver toxin to host cells^29^. While we demonstrated that purified ETEC OMV could bind directly to GM-1 gangliosides (supplemental figure 5B), we were unable to demonstrate substantial interaction between OMV and EV (supplemental figure 5c), and EV were ineffective in mitigating OMV-directed toxin delivery (supplemental figure 5D), suggesting that EV act primarily by engaging ETEC and free toxin.

**figure 4.**
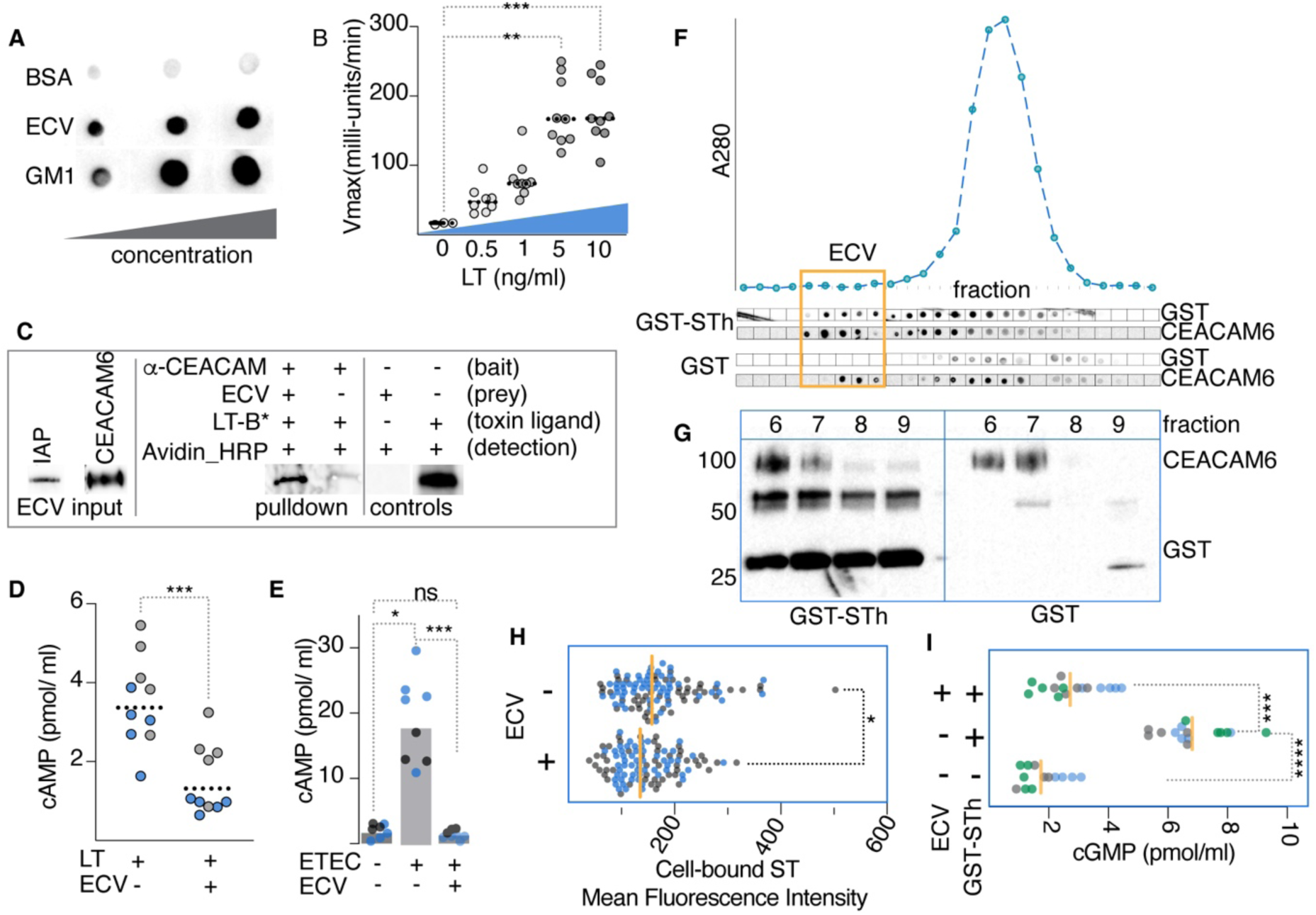
EV scavenge and neutralize ETEC toxins. **A.** EV contain ganglioside receptors for heat-labile toxin (LT). Shown are anti-LT dot immunoblots demonstrating LT-binding to increasing amounts of immobilized BSA (negative control, top) GM-1 ganglioside (positive control, bottom) and EV. **B**. Immobilized EV bind LT. Shown are kinetic ELISA in which EV bound to ELISA plates capture increasing amounts of LT. Summary of three independent experiments, ***0.0008, **0.0014 by Kruskal-Wallis comparisons to no LT control. **C**. Molecular pulldown study using anti-CEACAM antibody coated protein G beads (bait) to pull down EV (prey), and bound heat-labile toxin. Following incubation with biotinylated LT-B (LT-B*), immunoblot was developed with avidin-HRP to detect bound toxin subunit. Immunoblots (left panel) verify presence of intestinal alkaline phosphatase (IAP) and CEACAM6 in EV input prey. Biotinylated LT-B is indicated in blots of pulldown and controls. **D.** EV block LT-mediated activation of cAMP in target Caco-2 cells. Data reflect baseline-corrected values (raw data-baseline/baseline) and are from two independent experiments (n=10 total replicates). Analysis by Kruskal-Wallis. **E.** EV impede toxin delivery by ETEC. Caco-2 cAMP levels following infection with ETEC H10407 ± EV. Data are from two independent experiments (n=8 total replicates). Analysis by Kruskal-Wallis. **F.** Fractionation of GST-STh/EV complexes by size exclusion chromatography (SEC). Shown below the chromatogram are dot immunoblots for CEACAM6 and GST corresponding to individual fractions. Control fractions from GST-EV interactions are shown below. **G.** Western immunoblot of EV-containing fractions from SEC demonstrating co-elution of CEACAM6 and GST-STh. **H**. EV compete with T84 intestinal epithelial cells for ST binding. Shown are the results of two independent confocal microscopy experiments, with symbols representing mean fluorescence intensity of GST-STh binding in individual fields. Vertical lines represent geometric means. *p=0.02 (Mann-Whitney). **I.** EV neutralize STh activation of cGMP. Shown are results of 3 independent experiments (n=15 technical replicates). ****<0.0001, and ***n=0.0004 by Kruskal-Wallis.

In addition, we found that these same EV fractions also possessed guanylate cyclase C (GC-C), the receptor for heat-stable toxins (supplemental figure 5A). By size exclusion chromatography, we demonstrated that glutathione S transferase fused to STh (GST-STh) co-eluted with EV fractions containing CEACAMs (figure 4F<u>, G</u>), and that preincubation of the GST-STh fusion with EV impaired ST binding to target T84 cells (figure 4H), ultimately leading to significant reduction in toxin-mediated activation of cGMP (figure 4I). In summary these studies suggest that CEACAM-laden EV can engage ETEC and absorb both LT and ST effectively mitigating pathogen-host interactions by serving as molecular decoys for the bacteria as well as its secreted toxins.

### heat-labile toxin alters the composition of EVs

Although LT-mediated increases in cAMP, and subsequent activation of PKA are central to acute diarrhea caused by ETEC, PKA also governs the transcription of multiple host genes as it enters the nucleus to phosphorylate the cAMP-response element binding protein CREB^30,31^. Recent studies have demonstrated that LT modulates the transcription of multiple host genes in small intestinal epithelia ^17, 13,32^. To determine how the composition of EVs might be altered by LT we performed tandem mass spectrometry on vesicles isolated from 2D small intestinal enteroid monolayers with and without LT treatment (<u>supplemental dataset 1</u>). Interestingly, these studies also showed that the abundance of multiple proteins including both CEACAM6 and MUC2 were increased in abundance in EVs from LT-treated enteroids relative to those from controls (Figure 5, supplemental table 3). While the protein with the most increased abundance in LT-treated samples, FCGBP, which like MUC2 is also secreted by goblet cells, and intimately associated with mucin, its actual function remains undetermined^33,34^.

**figure 5.**
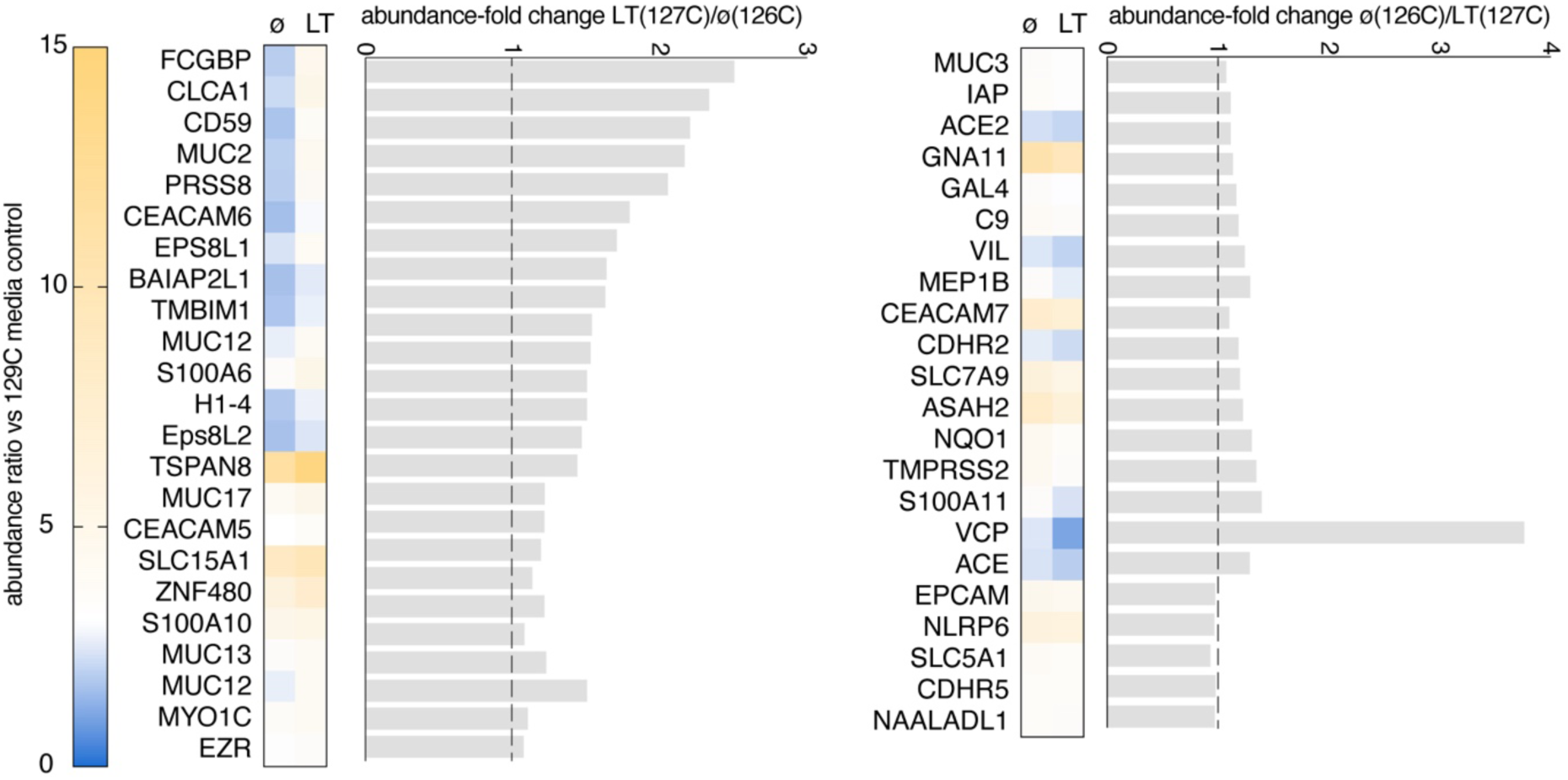
Comparative tandem mass tag spectrometry of EVs from LT-treated (127C label) and control (ø, 126C) enteroids. Subset of proteins identified in EVs: which were increased (left) and unchanged or decreased (right) following exposure of enteroids to LT (100 ng/ml overnight ∼18 hours).

We previously observed^32^ modulation of proteins linked to brush border biogenesis including the membrane adapter protein BAIAP2L1 involved in microvillus elongation^35,36^, or associated with exosomes including CD59 a membrane-bound complement regulatory protein^37,38^. Notably, our recent transcriptome studies of ileal enteroids also demonstrated that transcription of *myo1a*, a gene encoding microvillar motor protein ^26,39^ involved in EV biogenesis, is significantly depressed in following exposure to LT, leading us to question whether the quantity of EVs produced by epithelial cells would be impacted by LT exposure. However, short-term exposure (24 h) of small intestinal epithelia to LT had little appreciable impact on either the apparent size or quantity of EVs (supplemental figure 6A). Some suppression of EV production was observed with longer exposures to LT (72 h, supplemental figure 6B). Altogether however, toxin exposure appears to primarily drive changes in EV composition rather than the kinetics of EV biogenesis *in vitro*.

### ETEC infection enhances fecal shedding of CEACAMs

In CEACAM-expressing transgenic mice we observed increases in CEACAM shedding in feces following ETEC infection (supplemental figure 7). Examination of stools from children with ETEC diarrheal illness in Bangladesh demonstrated the presence of EV bearing the canonical EV marker CD9 and abundant CEACAMs (figure 6A). As previous immunohistochemistry studies of small intestinal biopsies obtained from ETEC-infected patients demonstrated that CEACAM6 production in the mucosa appeared to increase following infection^17^, and earlier studies also showed that CEA (CEACAM 5) is normally shed in significant amounts in human stool^18,19,40,41^, we questioned whether the shedding of CEACAMs also increased during human infection. To address this question, we developed a sandwich assay to capture CEACAM-laden material from fecal suspensions of ETEC-infected human hosts (figure 6B). We found that Bangladeshi patients with ETEC had appreciably higher amounts of CEACAMs in stool compared to healthy controls from the same endemic region(figure 6C), and that on challenge of human volunteers with ETEC, CEACAM content in stool transiently increases in the week following infection (figure 6D), further suggesting that expression of these molecules may play an important role in the innate response to ETEC infection in humans.

**Figure 6.**
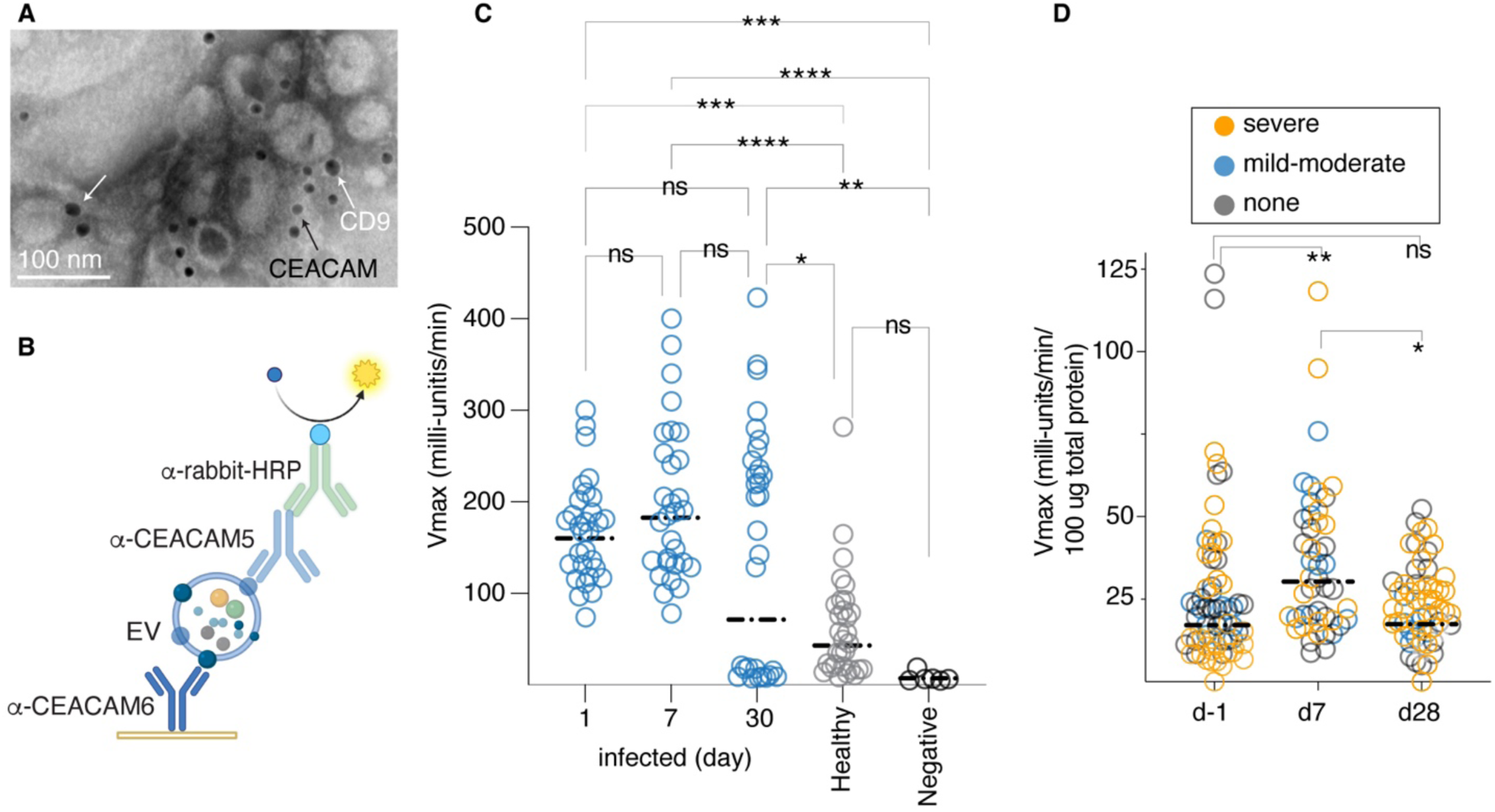
human ETEC infection is associated with increases in fecal CEACAMs. **A**. CEACAM-laden EV are shed in stool of children with acute ETEC diarrheal illness. Immunogold-labeled TEM image of EV isolated from diarrheal stool demonstrates detection of CD9 (larger 18 nm particles, white arrows), and CEACAMs (12 nm, black arrow). **B**. Schematic of kinetic ELISA strategy to capture and detect CEACAM+ EV from human stool.**. C**. Graph depicts summary of two independent experiments performed on samples from 5 patients naturally-infected with ETEC and 5 healthy controls (icddr,b in Dhaka), each with three technical replicates (total of n=30 data points for each day). Day 1 = day of presentation to icddr,b. Negative control wells contain only buffer used in sample extraction. **D**. Summary of two technical replicates in 2 separate experiments with samples obtained from human volunteers on the day prior to infection (d-1) and on days 7, 28 following challenge with ETEC H10407 (n=17; 3 with mild-moderate diarrhea, 8 with severe diarrhea and 6 with no diarrhea following challenge). Statistical analyses by Kruskal-Wallis nonparametric comparisons: ****p<0.0001, ***p=0.0002, **p=0.002, *p=0.0127.

## Discussion

Enterotoxigenic *E. coli* that cause infections in humans are largely host-restricted pathogens. Recently, we have demonstrated that ETEC engage gastrointestinal CEACAMs, in particular CEACAM6, to facilitate bacterial adhesion and toxin delivery to human intestinal epithelia^17^. Interestingly, the GPI-anchored CEACAMs including CEACAM6 are found exclusively in primates ^42^, potentially contributing to the unique relationship of ETEC to its human hosts. Indeed, CEACAM molecules appear to be at the center of an evolutionary “arms race” between pathogens and their human hosts. On the one hand, a diverse group of pathogens have evolved a variety of adhesins to engage these molecules as receptors. In contrast, under selective pressure of these pathogens, the host may deploy divergent or variant CEACAMs^43^ to minimize adhesin engagement and mitigate host tropism, or to target pathogens for destruction^43–46^.

The data presented here suggest a complex paradigm in which the same CEACAM is involved in bacterial adhesion and in innate host defenses. We previously demonstrated that the heat-labile toxin enhances bacterial adhesion by up-regulating target CEACAM6 on the surface of small intestinal enterocytes^17^. Here however, we show that LT also significantly increases CEACAM6 abundance in extracellular vesicles (EV) which can bind the bacteria at a distance to interdict effective adhesion to target enterocytes. Therefore, use of CEACAMs as receptors may ultimately come at some cost to the pathogen as they are eliminated by extracellular CEACAM-laden vesicles.

Notably, these EV also contain GM1 gangliosides, the established receptor for the B-subunits of both LT and the closely related cholera toxin, which like LT also upregulates production of CEACAMs on small intestinal epithelia^17^. Likewise, they bear GC-C, the receptor for heat-stable toxin (ST) as well as the endogenous peptides guanylin and uroguanylin. Notably, the affinity of ST for GC-C is 100 x that of guanylin and 10 x higher than uroguanylin, and both native peptides, unlike ST are subject to proteolysis^47^. Therefore, EV could aid in restoration of intestinal homeostasis by capturing the ST super agonist at a distance from receptors on the surface of epithelial cells, thereby eliminating competition with the lower affinity locally produced endogenous peptides. The ability to engage gastrointestinal pathogen(s) while effectively scavenging any secreted exotoxins may make EV a particularly effective element of innate intestinal defenses.

We found that other molecules potentially important in pathogen engagement were also concentrated in CEACAM-positive EVs. The Fc Gamma binding protein (<u>FCGBP</u>) most increased in abundance in EVs from LT treated enteroids, has previously been shown to be associated with exosomes^48,49^, and it has been suggested that these mucin-like molecules, which are concentrated in the intestinal goblet cells^50,51^, may play an important role in defense of the intestinal mucosa^52,53^. Similarly two other proteins shown here to be enhanced in EVs, CEACAM1 and CLCA1 are also associated with the mucin proteome^54–56^, findings that may relate to LT acting as a potent stimulus for goblet cell mucin production and secretion^57^.

The precise role of EV and the contribution of CEACAMs localized on their surface in the elimination of ETEC and other enteric pathogens deserves additional study. We should also note that in addition to the CEACAMs expressed on mucosal epithelia of the gastrointestinal tract, CEABAC10 mice also express low levels of CEACAM3 on neutrophils^23^. Studies of children with ETEC diarrhea have demonstrated significant amounts of lactoferrin as well as the leukocytes in stool^58^. In addition, single nucleotide polymorphisms in the lactoferrin gene have been associated with an increased risk of traveler’s diarrhea^59^, suggesting a role for recruitment of neutrophils in innate responses to ETEC infection. Although CEACAM6 and CEACAM3 share similar amino-terminal IgV-like domains^21^, our earlier studies suggested that ETEC preferentially engage CEACAM6 with little or no affinity for CEACAM3^17^. Overall, the current studies demonstrate that ETEC encounter CEACAM-laden vesicles *en route* to target intestinal epithelial cells, and that these have the potential to modulate the course of infection. The significant differences observed in the kinetics of infection between wild type and CEACAM-expressing transgenic mice may also point out limitations to the use of conventional mice in the investigation of ETEC and other *E. coli* pathovars.

Altogether, CEACAMs appear to play a bifunctional role in the pathogenesis of ETEC, acting both as toxin-induced receptors for these common pathogens as well as toxin-responsive molecular decoys for their elimination. These findings have important implications for the investigation of the molecular pathogenesis of ETEC and other gastrointestinal pathogens.

## Materials and Methods

### propagation of human small intestinal enteroids and transformed intestinal cells

Enteroids from human small intestine were propagated as previously described^17^. Briefly, cells originated from biopsy samples obtained from adults undergoing routine endoscopy with their consent and approval of the Washington University in Saint Louis School of Medicine Institutional Review Board. Stem cells derived from these samples were maintained in a biobank of the Precision Animal Models and Organoids Core (PAMOC) of the Washington University in Saint Louis Digestive Diseases Research Core Center (DDRCC).

Samples from human ileum (Hu235D) were re-suspended in Matrigel (BD Bisosciences, San Jose, California, USA), incubated at 37° C and 5% CO_2_ with 50% L-WRN conditioned media (CM) and 50% primary culture medium (Advanced DMEM/F12, Invitrogen) supplemented with 20% FBS, 2mM L-glutamine, 100 units/mL penicillin, 0.1mg/mL streptomycin, 10 µM Y-27632 (ROCK inhibitor, Tocris Bioscience, R&D systems, Minneapolis, MN, USA), and 10 µM SB431541 (TGFBR1 inhibitor, Tocris Bioscience, R&D systems).

After washing and trypsinization cells were centrifuged (1100 x g for 5 minutes), then re-suspended (1:1 CM and primary medium with Y-27632 and SB431541) as described above, and plated onto filters (Transwells®, 6.5 mm insert, 24 well plate, 0.4 μm polyester membrane, Corning Incorporated, Kennebunk, ME, USA) coated with collagen IV (Millipore Sigma). Inserts were rinsed with DMEM/F12 with HEPES, 10% FBS, L-glutamine, penicillin, and streptomycin, and cells grown to confluency in 50% conditioned media (CM). To differentiate monolayers media was changed to 5% CM in primary medium + ROCK inhibitor for 48 hours.

Caco-2 cells (ATCC <u>HTB-37</u>) were propagated at 37°C, in an atmosphere of 5% CO_2_, in Eagel’s Minimum Essential Media (MEM) supplemented with fetal bovine serum (FBS) to a final concentration of 20%. Cells were seeded and grown to confluence in 96 well plates for adhesion assays to determine bacterial adhesion by detergent lysis, or alternatively onto Transwell filters for confocal microscopy. T84 cells (ATCC <u>CCL-248</u>) were propagated in DMEM: F12 Media containing 5 % FBS.

### tandem mass tag-mass spectrometry (TMT-MS) of EVs

Sample proteolysis, isobaric mass tag labeling, peptide fractionation, and liquid chromatography-mass spectrometry of EVs isolated from small intestinal enteroids was conducted by the Mass Spectrometry & Proteomics Core, Johns Hopkins University School of Medicine. Samples were reduced, alkylated, and bound to SP3 beads for digestion with 25 µg/mL trypsin (Pierce, MSMS grade) in 100 mM triethylammonium bicarbonate (TEAB) for 16 h at 37 °C. Bead-bound peptides were then labeled in 100 mM TEAB with TMT Pro 16plex Isobaric Mass Tags (Pierce ThermoFisher). Peptides were fractionated by basic reverse phase chromatography and then analyzed on a nano-LC-Orbitrap-Lumos-ETD (ThermoFisher) interfaced with an EasyLC1000 series reverse-phase chromatography. Survey scans (full mass spectrum) were acquired within 375-1600 Da m/z using a Data Dependent Top 15 method with dynamic exclusion of 15 s. MS/MS spectra were searched with Mascot v.2.8.0 against the RefSeq2021_204_Human database. Search qualifiers included trypsin sites with missed cleavage 2 tolerance, precursor mass tolerance of 5 ppm, and fragment mass tolerance of 0.01 Da. Carbamidomethylation on Cys, TMT 16pro tag on N-terminus, and TMT 16pro tag on Lys were included as fixed peptide modifications. Oxidation on Met and deamidation of Asn and Gln were included as variable modifications. Peptide identifications from the Mascot searches were processed within Proteome Discoverer and Percolator to identify peptides with a confidence threshold of 1% False Discovery Rate, based on an auto-concatenated decoy database search, and to calculate the protein and peptide ratios. Only unique peptides were used for normalization and ratio calculations.

### CEACAM expressing transgenic mice

Transgenic mice which express multiple human CEACAM molecules (CEACAMs 3, 5, 6 and 7) were propagated from 2 female heterozygous CEABAC10 mice^23^ kindly supplied by the Gray-Owen laboratory at the University of Toronto. These were mated in the <u>Washington</u> <u>University Mouse Genetics Core</u> facility with C57BL/6N Crl (<u>Charles River Laboratories</u>) males. Pups generated from matings were genotyped to identify CEACAM+ heterozygotes.

Genotyping was performed on genomic mouse DNA extracted in hot sodium hydroxide/tris (HotSHOT)^60^. Briefly, 1-2 mm tail snips were dissolved in 75 μl of alkaline lysis buffer (25 mM NaOH, 0.2 mM disodium EDTA, pH 12) at 95°C for 30 minutes and then neutralized with equal volume of 40 mM Tris-HCl, pH 5. Presence or absence of the CEACAM5 gene was detected using primers-GACACAGCAAGCTACAAATGTGAAACCCAG (forward) and GCCACAGGTGATATTGTCAGAGGGAAGTGG (reverse) which amplify a 460bp amplicon. Both male and female mice were used throughout the studies for both CEABAC10 mice and C57BL/6N Crl controls.

### Intestinal challenge with enterotoxigenic *E. coli*

7-8 week-old mice were challenged with ETEC by orogastric lavage as previously described^61^. Briefly, mice were pretreated with streptomycin in drinking water (5 g/l) two days prior to challenge to reduce intestinal colonization with competing microbiota, and then returned to water without antibiotics 1 day prior. Two hours prior to challenge mice were treated with famotidine (1.25 mg in a volume of 125 µl) administered intraperitoneally (IP) to reduce gastric acidity, and fasted until gavage with ∼1.5 x 10^4^ colony forming units of jf876 (supplemental table 1). Stools were collected daily and fecal suspensions were diluted in PBS and plated onto Luria agar containing kanamycin (25 µg/ml). All studies in mice were conducted under protocol 20-0438 approved by the IACUC at Washington University in Saint Louis, School of Medicine.

### EV isolation and characterization

EVs were recovered from antibiotic-free supernatants of differentiated small intestinal enteroids propagated from human ileum (Hu235D). Supernatants were centrifuged at 1048 x g for 10 minutes to pellet debris and then concentrated ∼ 14-fold to a final volume of 0.5 ml (Amicon, 30K MWCO) prior to additional processing.

To isolate EVs, concentrated supernatants were separated by size exclusion chromatography (SEC) using resin with a 35 nm pore size (qEVoriginal/35 nm, Izon). The size distribution and quantity of EVs was determined by tunable resistance pulse sensing (TPRS) (qNano, Izon Ltd., Christchurch, New Zealand). All steps of nanopore optimization and sample measurement followed the guidelines outlined in the qNano Gold User Manual and Izon Control Suite software Custom Planner Tool (Izon). For the analysis, a nanopore NP100 (Pore ID A87921, Izon Ltd.) with an analysis range of 50−330 nm was utilized. The optimization of the nanopore was performed using polystyrene calibration particles (CPC100, Batch ID 20221003, Izon Ltd.) with an average particle diameter of 100 nm and a concentration of 1.8E+13 for calibration purposes. Both the calibration particles and the samples were run under the same conditions, including stretch, pressure, voltage, and baseline current. To eliminate the impact of pore and particle charge on the detected concentration, all samples were analyzed at two pressure points.

### Dot-immunoblotting of SEC fractions

2 µl of each fraction was spotted onto nitrocellulose membranes, dried at 37°C for 5 minutes, blocked with 5% milk in PBS, 0.05% Tween-20 for 30 minutes at 37°C, then incubated with primary antibodies against CEACAM6 (9A6), CD9 (C-4), lysozyme, or intestinal alkaline phosphatase diluted 1:1000 in 2.5% milk in PBS, 0.05% Tween-20 for 1 hour at 37°C. After washing 3x in PBS, membranes were incubated with the respective anti-mouse or anti-rabbit HRP-conjugated secondary antibodies (1:1000 dilution in PBS) for 45 minutes at room temperature, washed again in PBS, and developed with Clarity ECL Western blot substrate (Bio-Rad,<u>1705061</u>). To detect binding of the cholera toxin B subunit, fluorophore-conjugated CT-B (ThermoFisher <u>C34775</u>) was incubated with membranes prepared as above at a final concentration of 4 µg/ml in PBS 0.05% Tween-20 for 30 minutes at room temperature, washed 3 x im PBS and imaged on an Azure biosytems c600 molecular imager.

To detect CEACAMs in the feces of CEABAC10 mice following ETEC infection, fecal pellets were collected prior to infection, and on days 1 and 9 post infection. 300 μg from each mouse was resuspended in resuspension buffer (PBS-0.5% Tween-20 containing 5 mM sodium azide) by vigorous vortexing and centrifuged at 845 x g for 30 minutes at 4°C. Clarified supernatant (2 µl) was dotted onto nitrocellulose as above and probed with rabbit polyclonal anti-CEA primary antibody (1:1000; Dako, Denmark A0115) and HRP conjugated anti-rabbit secondary antibody (1:1000; Rockland, 611-1322). Blots were visualized by chemiluminescence using Clarity Western ECL substrate (Bio-Rad <u>1705061</u>) and signal intensities were measured using ImageJ2 v2.14.

To isolate extracellular vesicles from CEABAC10 transgenic mice and littermate controls, mice were sacrificed, the small intestine excised, and flushed 3 times with 5 ml of PBS supplemented with protease inhibitor (Pierce Protease Inhibitor Mini, Thermo Scientific). Debris was removed from the lavage by centrifugation at 1048 x g for 5 minutes, followed by passage of lavage fluid through a 70 µm filter, and concentration through a 3K MWCO filter (Amicon Ultra-4). EVs were then isolated by size exclusion (qEVoriginal/35 nm, Izon).

To isolate extracellular vesicles from human fecal specimens, diarrheal samples from ETEC infected patients (n=5) were pooled together and resuspended in 15 ml PBS supplemented with protease inhibitors (Pierce Protease Inhibitor Mini, ThermoFisher Scientific, A32955), centrifuged at 4°C at 875 x g, filtered through 70 µm filter and concentrated using a 30 kDa MWCO filter (Amicon Ultra-15, Millipore).

Total protein concentration of each vesicle preparation was assessed using the Qubit Protein Assay Kit (<u>Q33212</u>, ThermoFisher). To ensure complete lysis, vesicle samples were incubated with 0.2% SDS for 10 minutes at 95°C. Samples and standards were then incubated with Qubit working solutions for 15 minutes at room temperature and read with a Qubit Fluorometer 3.0.

In CEACAM depletion experiments, anti-CEA rabbit polyclonal antibodies (supplemental table 2) were immobilized onto protein G Dynabeads (Invitrogen) and incubated with culture supernatant at room temperature for 1 hour. CEACAM-depleted supernatant was then separated from beads by magnetic separation. CEACAM depletion was verified by immunoblotting.

### transmission electron microscopy

Transmission electron microscopy (TEM) of small intestinal enteroids infected with ETEC was performed in the <u>Department of Molecular Microbiology Imaging Facility.</u> After gentle washing with PBS, samples were first fixed in a solution of 2% paraformaldehyde/2.5% glutaraldehyde (Ted Pella, Inc., Redding, CA) in 100 mM sodium cacodylate buffer, pH 7.2 for 2 hours at room temperature. Samples were then placed at 4°C overnight, and then washed in sodium cacodylate buffer and postfixed in 2% osmium tetroxide (Ted Pella, Inc) for 1 hour at room temperature. After rinsing in deionized water, samples were dehydrated in ethanol, and embedded in Eponate 12 resin (Ted Pella, Inc.) cut into sections (95 nm) with an ultramicrotome (Leica Ultracut UCT, Leica Microsystems, Inc., Bannockburn, IL), and stained with uranyl acetate and lead citrate.

To immunolabel vesicles, fractions were absorbed onto glow-discharged formvar/carbon-coated nickle grids (<u>Ted Pella, Inc</u>.) for 10 min followed by negative staining. Grids were then washed with PBS and blocked with 1% FBS for 5 min Grids were subsequently incubated with rabbit anti-CEA (<u>Dako,</u> supplemental table 2) for 30 min. Grids were then incubated with secondary goat anti-rabbit IgG antibody conjugated to 12 nm colloidal gold (Jackson ImmunoResearch Laboratories, Inc. <u>111-205-144</u>, West Grove PA) for 30 min. Grids were then washed, fixed with 1% glutaraldehyde, and stained with 1% aqueous uranyl acetate (Ted Pella Inc., Redding CA) for 1 min. Excess liquid was gently wicked off and grids were allowed to air dry. Samples were viewed on a JEOL 1200 EX transmission electron microscope (JEOL USA Inc., Peabody, MA) equipped with an AMT 8 megapixel digital camera and AMT Image Capture Engine V602 software (Advanced Microscopy Techniques, Woburn, MA).

Intestinal lavage specimens from CEABAC10 mice challenged with ETEC strain jf876 (serotype O78) (table S1) were processed by SEC as above, and concentrated material was then used to identify EV-coated bacteria by immunogold TEM. Grids were incubated with rabbit anti-O78 antisera (Penn State) followed by anti-rabbit 18 nm gold conjugate (Jackson ImmunoResearch) to identify ETEC and mouse anti-CEACAM6 monoclonal antibody (9A6, Santa Cruz) followed by anti-mouse 12 nm gold conjugate.

### Purification of recombinant GST-STh fusion protein

Recombinant glutathione S transferase (GST) and GST-STh fusion proteins were purified as previously described^62^. Briefly bacterial strains jf1364 and jf3265 were grown overnight at 37°C, 225 rpm from frozen glycerol stocks in 2 ml of Luria Broth containing carbenicillin (100 µg/ml), diluted in fresh media and grown to OD600 of ∼0.6 then induced with isopropyl-β-d-thiogalactopyranoside (IPTG) for 2 hours. Cell pellets were extracted by sonication 5x in the presence of protease inhibitor (Roche Complete <u>11697498001</u>). Clarified supernatants were loaded onto 3 ml columns packed with glutathione agarose resin (GoldBio <u>G-250-10</u>), and after washing with PBS, recombinant GST or GST-STh was eluted with buffer containing 100 mM Tris-HCl, pH 8.0 and 10 mM reduced glutathione (MilliporeSigma 70-18-8), then dialyzed vs PBS.

### ETEC adhesion-inhibition assays

ETEC strain H10407 (supplementary table 1) was grown from frozen glycerol stocks in LB media at 37°C under static conditions as previously described^16^ to enhance expression of type 1 pili. Bacteria were added to Caco-2 cells to achieve a MOI (multiplicity of infection) of ∼1:10 and incubated with either CEACAM-enriched concentrated supernatants, CEACAM-depleted supernatant, or PBS as a control. To assess the effects of extracellular vesicles (EV) isolated from culture supernatants, the inoculum was incubated with vesicles for 15 minutes at 37°C prior to infection. Following inoculation, polarized Caco-2 cell monolayers were incubated at 37°C in a humidified tissue culture incubator with 5% CO_2_. After incubation for an hour, the cell monolayers were washed three times with gentle shaking (100 rpm on an orbital shaker for 1 minute per wash) using pre-warmed media to remove any unbound bacteria. Cell monolayers were then lysed with 0.1% Triton X-100 for 5 minutes, and the lysates were plated on LB-agar and grown overnight at 37°C to enumerate colony forming units (CFU). Alternatively, infected monolayers on Transwell filters were fixed with 4% PFA for 30 minutes at 37°C prior to immunofluorescence staining.

### Toxin binding

EV immobilized on nitrocellulose membranes, and blocked with 5% milk in PBS containing 0.05% Tween-20, were probed with double mutant heat-labile toxin (dmLT, L192G/L211A) at a concentration of 4 µg/ml to examine binding of LT to EV. LT binding was detected with mouse antisera raised against dmLT^63^ (1:1000), followed by horse anti-mouse IgG conjugated to HRP (Cell Signaling <u>7076</u>, 1:1000), and developed with ECL substrate. Blots were then imaged on an Azure biosytems c600 molecular imager. Alternatively, EV (0.22 mg/ml in a final volume of 100 µl/well) were immobilized on ELISA plates (Costar, 2580) incubation overnight at 4°C. The following day plates were washed and blocked with 5%BSA in PBS for 1 hour at 37°C. After washing with PBS, plates were incubated with increasing concentrations of LT for 1 hour at 37°C. After washing 3x with PBS, bound LT was detected using mouse polyclonal antisera against LT (1:1000, x 1 hour at 37°C), washing 3x with PBS, incubated with HRP-conjugated horse-anti-mouse IgG secondary antibody (Cell Signaling <u>7076</u>; 1:2000, x 1 hour at 37°C), and developed with 3, 3’, 5, 5’ - tetramethylbenzidine peroxidase substrate (TMB, sera care <u>5120-0053</u>). ELISA readings were acquired kinetically and recorded as Vmax (milli-units/min) (Eon, BioTek).

### EV-toxin Molecular interaction assays

Purified heat-labile toxin B subunit (LT-B), graciously provided by John D. Clements, Tulane University, was biotinylated (EZ-Link Sulfo-NHS-LC-Biotin) (Thermo Scientific <u>21335</u>) according to the manufacturer’s protocol, and dialyzed to remove excess biotin. Biotinylated LT-B ligand (10 µg) was then added to purified EV (∼11 µg) and incubated for 1 hour at 37°C in a final volume of 100 µl. An equal volume of PBS containing biotinylated LT-B was used as a negative control. Protein G magnetic beads (<u>Invitrogen 10003D</u>) were combined with anti-CEA antibody (Dako), and then used to capture EV for 1 hour at room temperature. Following magnetic separation, beads were washed in PBS then incubated in SDS loading dye for 15 minutes at 95°C. Solubilized proteins were resolved by 10% SDS-PAGE, and transferred to nitrocellulose then developed with Avidin-HRP (BioRad <u>1706528, 1:25,000</u>) followed by enhanced chemiluminescent (ECL) substrate (ThermoFisher Scientific <u>34094</u>).

Size exclusion chromatography was performed to demonstrate interaction between heat-stable toxin (STh) expressed as a recombinant GST fusion protein and EV. Supernatant media from Hu235D small intestinal enteroids was first concentrated (∼7-fold) through a 30 kD MWCO filter (Amicon Ultra-15, Millipore Sigma <u>UFC903024</u>) to a final volume of ∼1 ml. 500 µl of concentrate was then incubated for 30 minutes at 37°C with either GST alone or GST-STh. Mixtures were then subjected to size exclusion chromatography (<u>35 nm qEVoriginal</u>, Izon), fractions collected and saved at -80°C for subsequent analysis by dot immunoblotting with antibodies against GST (Invitrogen 13-6700), or anti-CEACAM6 (Santa Cruz 9A6), followed by anti-mouse IgG HRP conjugated antibodies (Cell Signaling 7076S). Fractions 6 to 9 which showed maximum GST and CEACAM6 signals in the dotblot assay were resolved by 10% SDS-PAGE, and Western immunoblots processed as above and developed with enhanced chemilumincent substrate (ThermoFisher <u>34094</u>).

### Toxin neutralization by EV

To investigate the impact of EV on ETEC toxin delivery target Caco-2 cells (ATCC <u>HTB-37</u>) were seeded in 96 well tissue culture plates at a density of ∼3 x 10^4^ cells/well, and incubated at 37°C, 5% CO_2_ for 48 hours. H10407 was grown under static conditions at 37°C as previously described^16^, and ∼ 10^6^ cfu (Multiplicity of infection∼100:1) were added per well with competing EV (∼11.5 µg/well) an equal volume of media. Following addition of bacteria ± EV, infected monolayers were incubated at 37°C, 5 % CO_2_ for 2 hours, washed to remove excess bacteria, media replaced, and then cellular cAMP was determined by competitive ELISA (Arbor Assays, <u>k019-h</u>). To examine the ability of EV to bind and neutralize LT, 2.5 ng of LT was added to 11.5 µg of purified EV or an equivalent volume of PBS. After incubation for 1 hour at 37°C, the LT ± EV mixtures were added to target Caco-2 cells and incubated for 18 hours at 37°C, 5 % CO_2_ prior to determination of cAMP levels as above. For ST neutralization, 1 mg of GST-STh in a volume of 1 ml was reduced with DTT (5 mM) for 3 hours at room temperature. After addition of 2 units of native bovine protein disulfide isomerase (PDI) (Creative Enzymes, <u>NATE-0533)</u>, the sample was dialyzed overnight against 1 liter of PBS. Following incubation with EV or buffer control, samples were passed through a 100K MWCO filter (Amicon <u>UFC510008</u>) to retain bound GST-STh. Filtrates were diluted 2 fold in tissue culture media then added to target T84 cells in the presence of phosphodiesterase inhibitors (25 µM), incubated for 4 hours at 37°C, 5% CO_2_, and intracellular cGMP determined (Arbor Assays <u>K065</u>).

To detect binding of GST-STh to T84 cells, ∼ 50,000 cells were added to Transwell filters (Costar 3470) and propagated for 5 days in DMEM/F12 supplemented with 5% FBS. After washing with 3 x with PBS, cells were incubated for 30 minutes at 37°C with 10 ng/ml of GST-STh in PBS ± 50 µl of Hu235D EV (220 µg/ml total protein concentration), then washed 3 x with PBS. After fixation with 4% paraformaldehyde (37°C x 10 minutes, room temperature x 20 minutes), filters were washed with PBS, then incubated with 2% BSA in PBS for 30 minutes at room temperature. GST-STh was detected with GST-cross-absorbed rabbit anti-GST-STh antibodies^62^ followed by goat-anti-rabbit IgG AlexaFluor 488 conjugated secondary antibody. Immunofluorescence signals in confocal images were then quantified with NIS-Elements AR software (Nikon 5.11.01).

### Confocal microscopy of cell associated bacteria

Cell-associated bacteria were detected using anti-O78 rabbit primary antibody, followed by Alexa Fluor 488 fluorophore conjugated goat anti-rabbit secondary antibody (<u>A11008</u>, ThermoFisher) and imaged by confocal microscopy (Nikon ECLIPSE Ti2). Nuclei were stained with 4′,6-diamidino-2-phenylindole dihydrochloride (DAPI, Sigma). Images were processed using NIS-Elements AR software version 5.11.01,(Nikon).

### Myeloperoxidase Immunohistochemistry of ETEC-infected CEABAC10 intestinal tissue

Small intestinal tissue sections cut from formalin-fixed paraffin-embedded blocks were mounted onto glass slides, then deparaffinized with xylene and treated with 3 % H_2_O_2_ in methanol for 15 minutes. Antigen unmasking was performed using heat retrieval in Diva Decloaker (Biocare Medical <u>DV200MX</u>) in a pressure cooker at 15 PSI and 99°C for 3 minutes. The sections were then blocked with 1% BSA, 10% normal goat serum in PBS. Slides were incubated with anti-MPO antibody (ab20670, Abcam) at a 1:500 dilution overnight at 4°C, washed, and then incubated with VisuCyte Rabbit HRP Polymer (VC003-025, R&D) at 1:4 for 1 hour at room temperature. After washing, the slides were developed with DAB (3,3′-diaminobenzidine, Vector SK-4100) as per the manufacturer’s protocol, washed, and counterstained with hematoxylin. Brightfield images were obtained with a BZ-X810 microscope (Keyence, IL).

## Flow cytometry

### Flow cytometry analysis of CEACAM-coated bacteria

We infected human CEACAM-expressing CEABAC10+ adult mice with GFP-expressing ETEC. Two days post-infection, fecal pellets were collected. One hundred micrograms (100 µg) of fecal pellets were resuspended in 1 ml of PBS and kept on ice for 15 minutes to allow debris to settle. The liquid portion was collected from the top and centrifuged at 3381 x g for 5 minutes to pellet down bacteria. To detect CEACAM-coated bacteria, pellets were then stained with anti-CEA primary antibody (DAKO) at a 1:200 dilution for 1 hour on ice, washed three times with PBS, and then incubated with goat anti-rabbit Alexa-Fluor 594-conjugated secondary antibody at a 1:200 dilution for 1 hour. For each sample, a tube without the anti-CEA primary antibody, but with the secondary antibody, was used as an unstained control. All samples were stained with DAPI at a 1:1000 dilution prior to acquisition. Samples were analyzed using FlowJo software (version 10.9.0). GFP+DAPI+ double-positive events were gated for ETEC, from which the amount of CEACAM-positive and CEACAM-negative bacteria was determined based on the corresponding unstained control.

### Live-dead staining

To assess the bactericidal activity of membrane vesicles, we conducted live-dead staining followed by flow cytometry analysis. Both log-phase and stationary phase ETEC cultures were grown in the presence or absence of vesicles for 1 hour, 2.5 hours, and 3.5 hours under different conditions (static or shaking) and in different media (LB, cell culture media, or PBS). After the incubation, bacterial samples were pelleted and resuspended in a live-dead staining buffer (PBS supplemented with 1mM EDTA and 0.1% Na-azide, pH 7.4). To each sample, a final concentration of 50 µM propidium iodide and 420 nM thiazole orange dye was added, followed by vortex mixing and a 5-minute incubation at room temperature. Samples were acquired using a BD FACSCalibur and analyzed with FlowJo software.

### immunodetection of CEACAMs in stool

Stool specimens obtained from patients at icddr,b during natural ETEC infection, healthy controls, or from earlier ETEC controlled human infection model studies^64^ were shipped on dry ice and maintained at -80°C prior to use. ELISA wells (Costar EIA <u>2580</u> Corning, Kennebunk, ME, USA) were coated overnight (4°C) with 100 µl/well of CEACAM6-specific monoclonal antibody (9A6) diluted 1:100 in 50 mM carbonate buffer, pH 9.6. Plates were then washed 6 times with 1x PBS (pH 7.4, Corning) containing 0.05% Tween-20 (Sigma), and blocked with 1% BSA in PBS for one hour at 37°C. Stool samples were extracted in PBS-0.5% Tween-20 containing 5 mM sodium azide, and centrifuged at 845 x g for 30 minutes at 4°C. Clarified supernatants were then diluted 1:10 in PBS. 30 µl of each sample was added per well, and incubated at 37°C for one hour, after which plates were washed 6x with PBS-0.05 % Tween. To detect bound CEACAMs 100 µl of polyclonal rabbit anti-CEA antibody (Dako, Denmark A0115) diluted 1:1000 in PBS, 0.5% BSA was added per well and incubated at 37°C x 1 hour. Plates were again washed 6x with PBS-Tween, and then incubated with 100 µl of HRP-conjugated goat IgG anti rabbit IgG (H&L) (Rockland, 611-1322).

All human studies were approved by the Institutional Review Board at Washington University in Saint Louis School of Medicine under protocol number 201110126.

### Data sharing

Mass spectrometry and original image data are available on Figshare with Digital Object identifiers in supplemental table 4.

### Statistical analyses

Mann-Whitney was used to compare two unpaired groups of nonparametric data. Kruskal-Wallis was used in comparison of three or more groups of data with Dunn’s multiple comparisons test. The log-rank (Mantel-Cox) test was used in comparison of survival curves.

## acknowledgements

JMF was supported by funding from the Department of Veterans Affairs (5I01BX001469-05), and the National Institute of Allergy and Infectious Diseases (NIAID) of the National Institutes of Health (NIH) R01 AI170949, R01 AI089894, and support by the NIH Washington University DDRCC Grant NIDDK P30 DK052574. Research conducted by AS was also supported by National Institute of Allergy and Infectious Diseases of the National Institutes of Health under Award Number T32AI007172. Mass Spectrometry was conducted in the Proteomics Core of the Johns Hopkins Conte Digestive Diseases Basic and Translational Research Core Center (P30 DK-089502) with the assistance of the Proteomics Core Director, Robert Cole, Ph.D. The content is solely the responsibility of the authors and does not necessarily represent the official views of the National Institutes of Health, or the Department of Veterans Affairs.

