## supplemental information for "Host-derived CEACAM-laden vesicles engage enterotoxigenic *E. coli* for elimination and toxin neutralization"

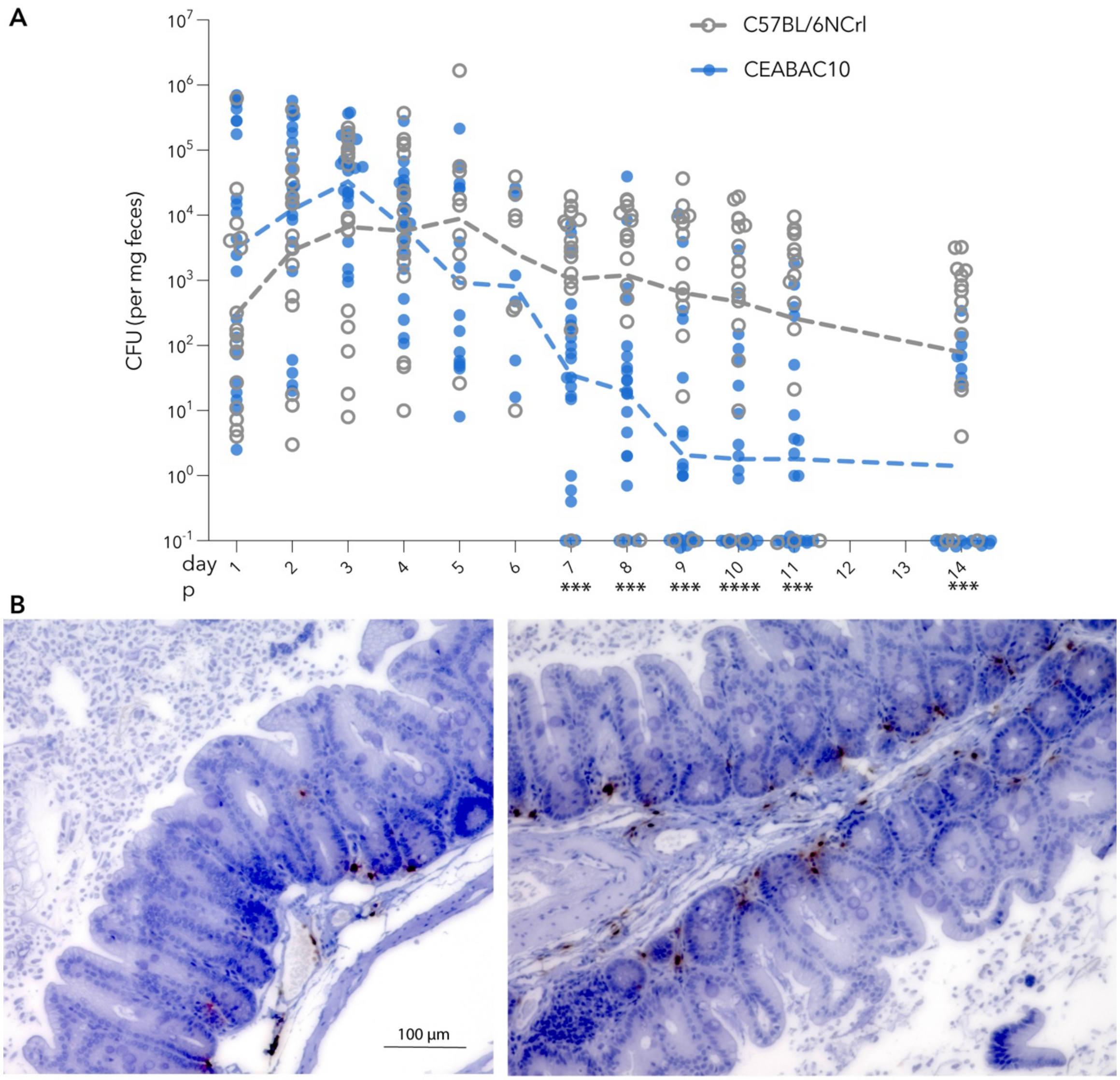


**Fig. S1. CEACAM expression enhances ETEC elimination independent of neutrophil infiltration. A.** Shown are data combined two independent colonization experiments with a total of n=21 CEABAC10, n=17 C57BL/6NCrl (controls). Dashed lines connect geometric means. ***<0.001, ****<0.0001 by Mann Whitney two-tailed nonparametric comparison between groups. **B.** Myeloperoxidase staining is confined to basolateral surface of the small intestine of ETEC-infected CEABAC10 mice. Shown are two immunohistochemistry sections of ileum from ETEC-infected CEABAC10 mice showing the distribution of myeloperoxidase activity (associated with neutrophils) in basolateral regions. Brown color indicates detection by DAB (3,3'-diaminobenzidine) oxidation.

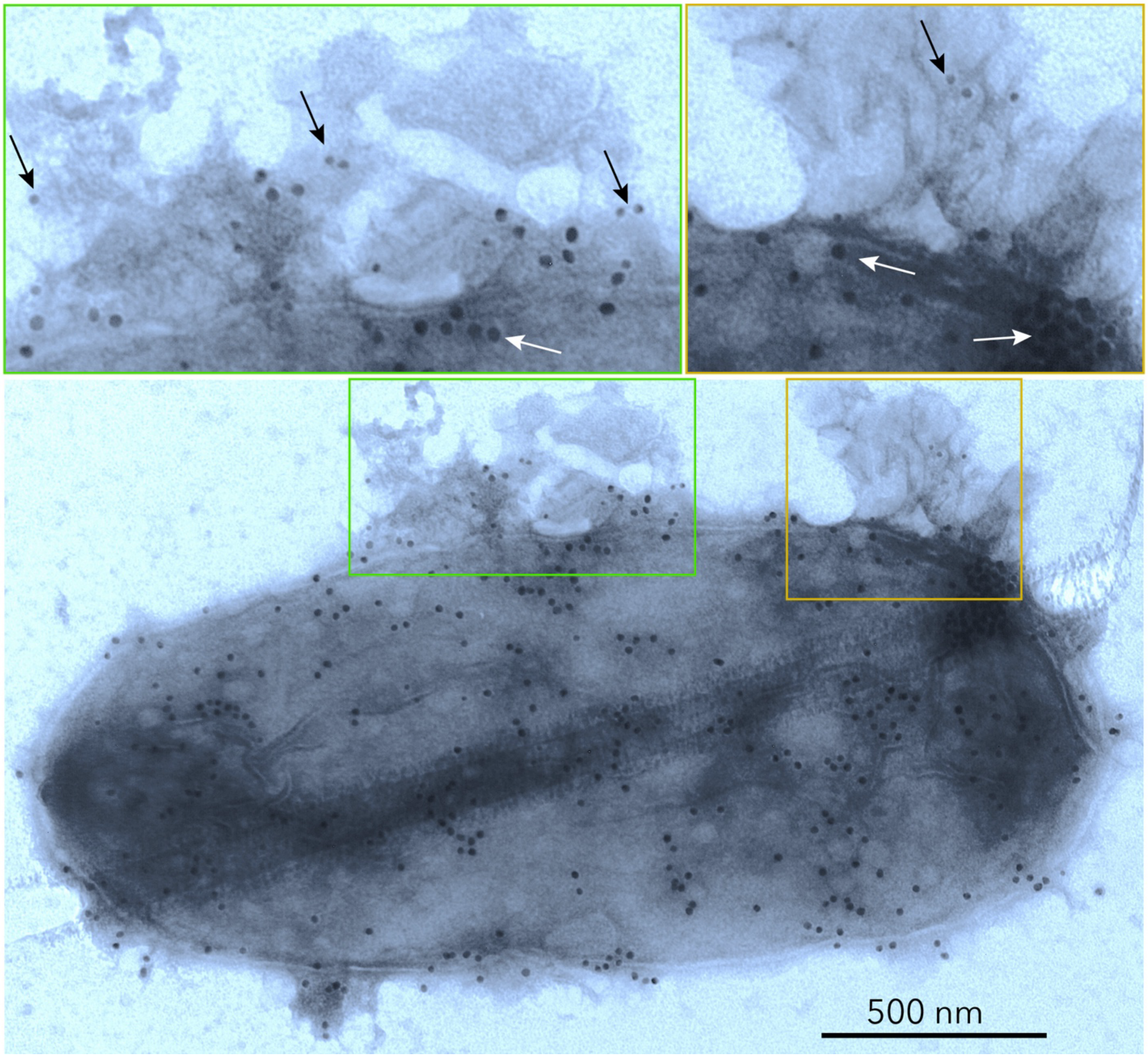


**Fig S2. ETEC coated with CEACAM+EV emerge in stool following challenge.** Immunogold transmission electron micrograph shows ETEC identified in fecal extracts of CEABAC-10 mouse following challenge with H10407 (serotype O78). Larger 18 nm gold particles on the bacterial surface (white arrows, insets) were used to identify O78 oligosaccharide and while CEACAM6 was labeled with smaller 12 nm gold (black arrows, insets).

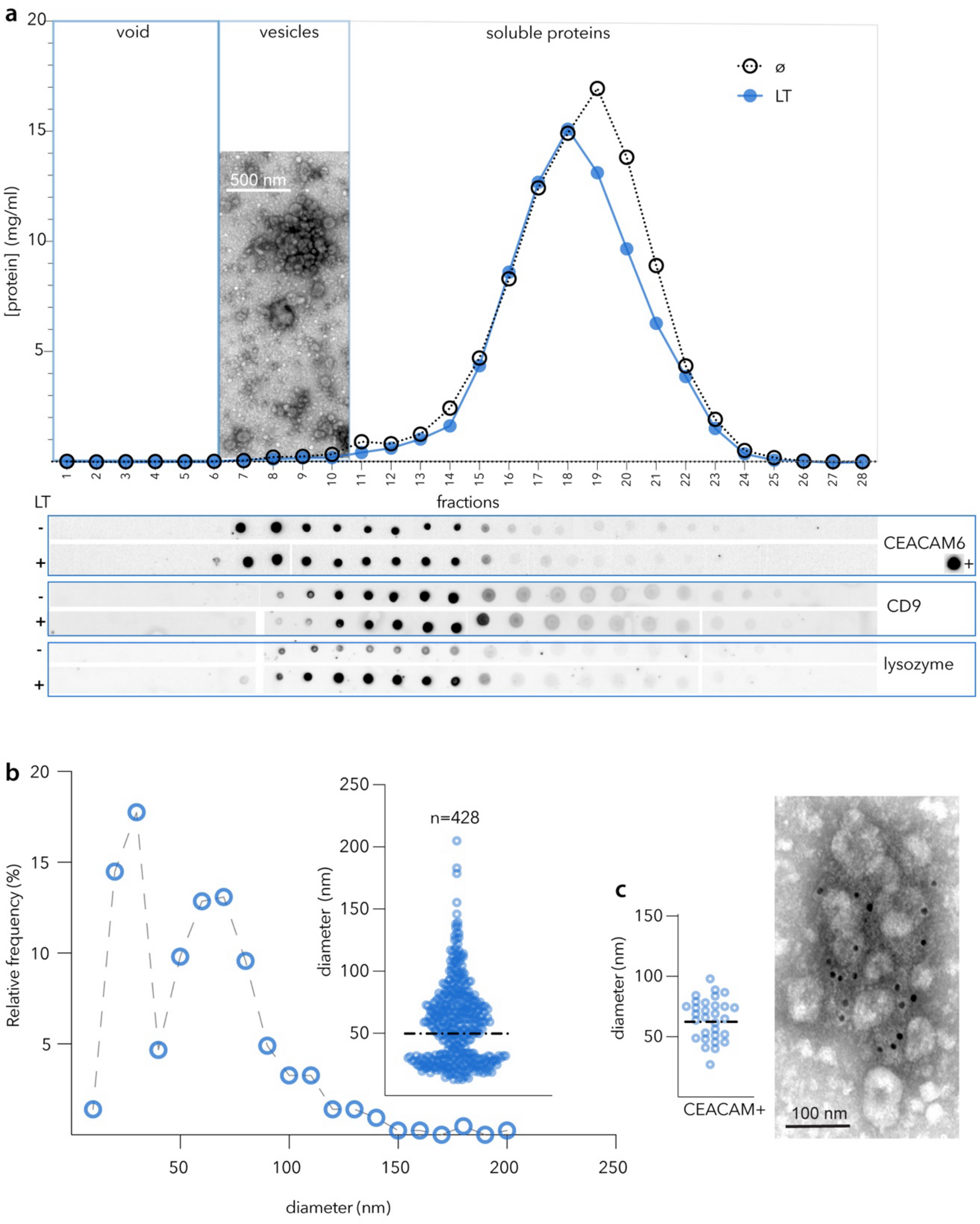


**Fig S3**. isolation and characterization of extracellular vesicles (EV) from small intestinal enteroid supernatants. **a**. chromatogram of EV isolation from LT-treated and untreated (ø) enteroids by size exclusion chromatography (Izon, qEV 70 nm). Dot immunoblots below indicate CEACAM6+, CD9, and lysozyme+ fractions. Inset EM shows EV pooled from fractions 6-10. **b**. size distribution of isolated vesicles (dashed horizontal line =geometric mean diameter). **c.** size distribution of CEACAM + extracellular vesicles (representative immunogold EM at right).

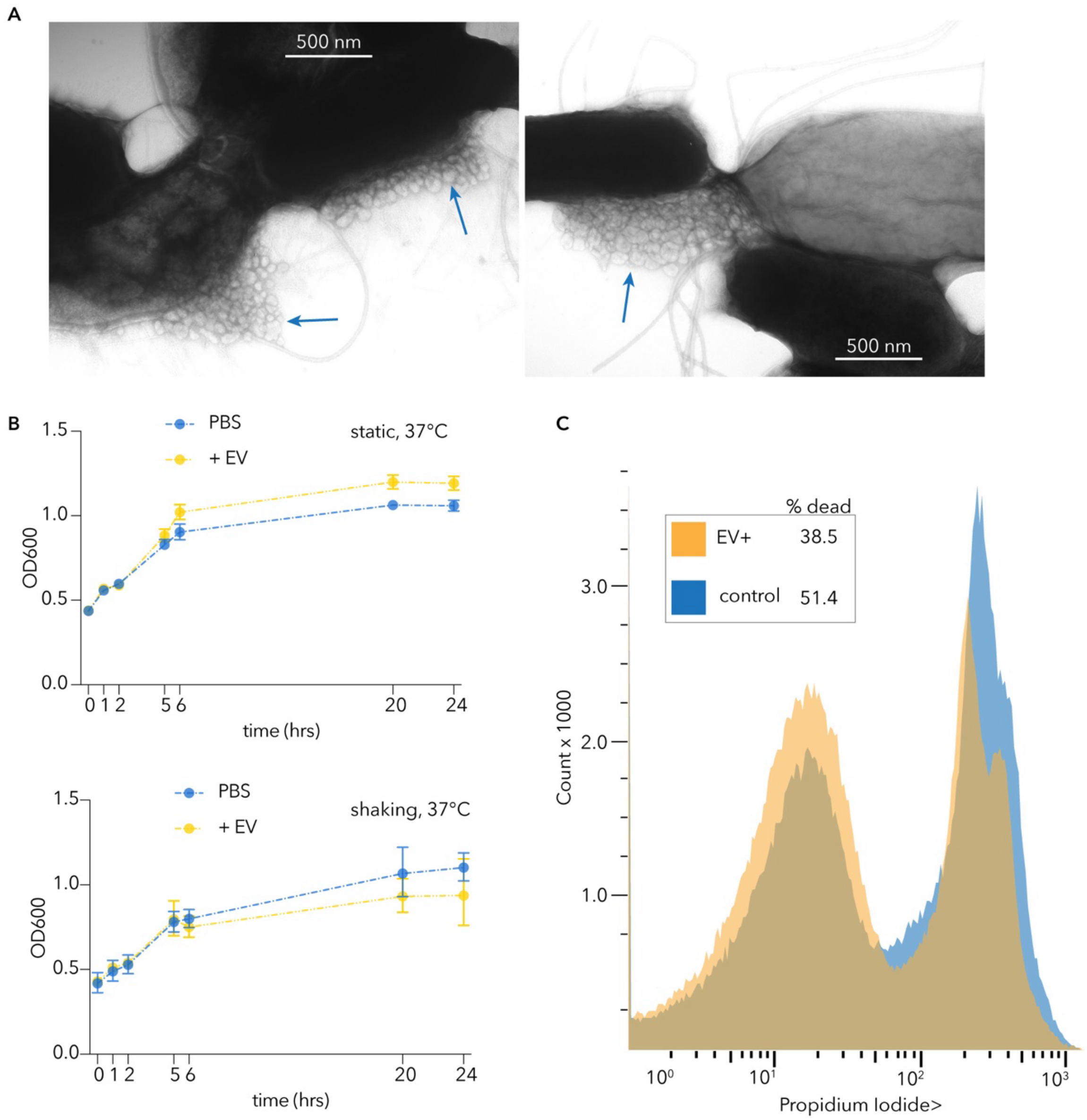


**Fig. S4.** Purified host-derived extracellular vesicles (EV) bind but do not kill ETEC. A. purified extracellular vesicles (arrows) bind to the surface of ETEC. B. growth curves of ETEC ± EV (C). live-dead staining of ETEC ± EV analyzed by flow cytometry.

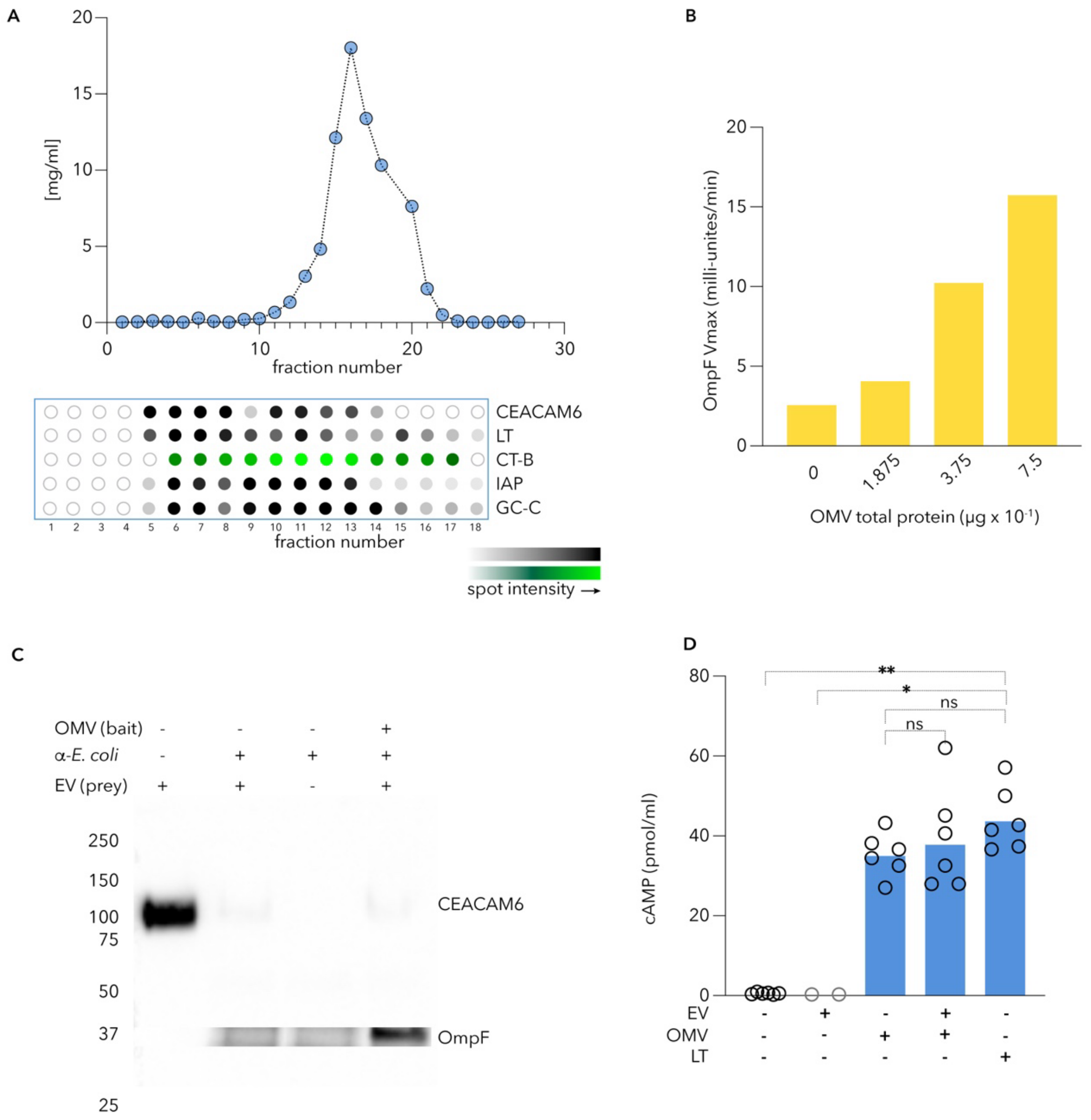


**Fig. S5. CEACAM6+ EV contain enterotoxin receptors**

**A.** Graph represents protein concentration of individual size exclusion chromatography fractions. Shown below are immuno dot-blot images (gray-scale) for indicated proteins (right) present in SEC fractions and binding of fluorescently-labeled CT-B (green) to each fraction. Fraction numbers 1-18 (bottom). **B**. ETEC outer membrane vesicles (OMV) bind to GM-1 gangliosides (shown are kinetic ELISA data demonstrating detection of OmpF following OMV binding to target GM-1 gangliosides N=5 technical replicates per concentration. **C**. Molecular pulldown studies with ETEC OMV (bait) and CEACAM6+ EV (prey), demonstrate minimal interaction between bacterial and host vesicles. **D.** OMV deliver toxin to target Caco-2 cells but are not blocked by EV. Shown are cAMP levels following exposure to EV alone; ETEC H10407 OMV; OMV added in the presence of host EV (isolated from Hu235D); and LT alone (+ control). **=0.006, *=0.04, ns=nonsignificant (Kruskal-Wallis).

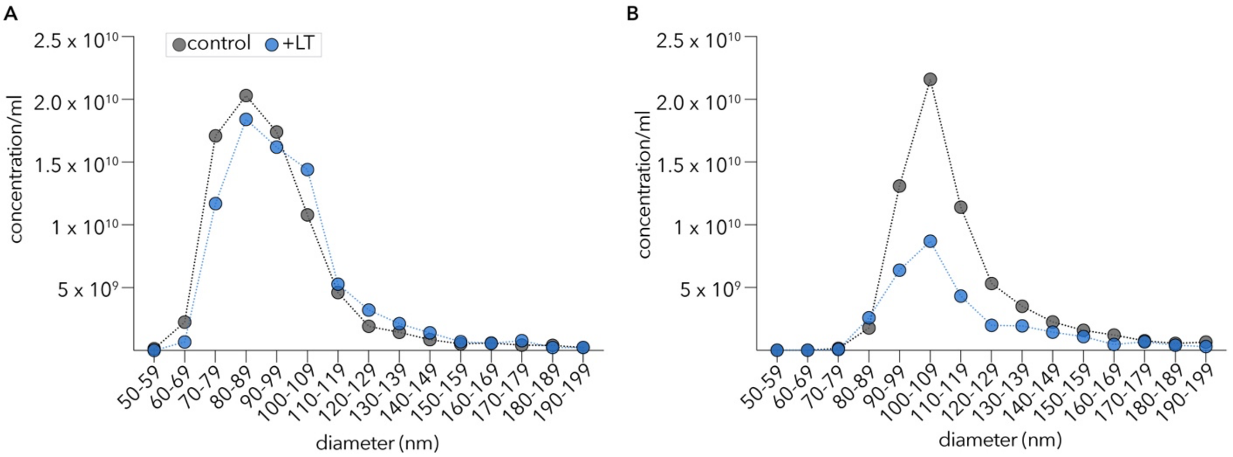


**Fig. S6.** Profiles of EV isolated from control untreated enteroids and LT-treated cells at (A) 24 and (B) 72 hours.

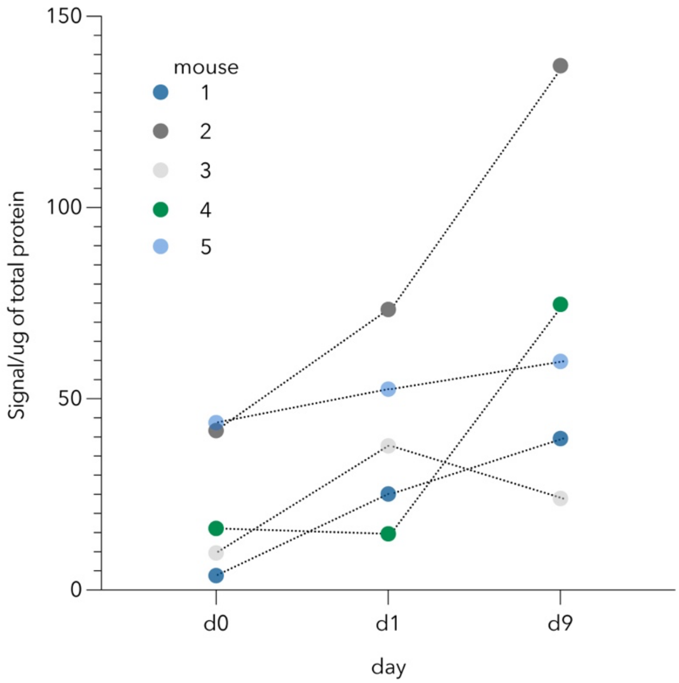


**Fig. S7.** Fecal CEACAMs increase in CEABAC10 mice following challenge with ETEC. Shown are results of immuno-dot blot screening of fecal suspensions for CEACAMs in n=5 CEABAC10 mice at baseline (d0) and following infection (day 1, and day 9) with ETEC H10407.

### Table S1. bacterial strains and plasmids used in these studies

| **supplemental table 1 bacterial strains and plasmids used in these studies** | | |
|---|---|---|
| strain | description | reference(s) |
| H10407 | wild type ETEC strain, *eltAB*, *estP, estH;* Serotype O78:H11 | (1) |
| jf876 | *lacZYA*::$Km^R$ derivative of H10407 | (2) |
| jf1364 | BL21(DE3) (pGEX-3X) | |
| jf2450 | H10407(pGFPmut3.1), $Amp^R$ | (3) |
| jf3265 | Top10(pCW002) | (4) |
| plasmids | | |
| pCW002 | pGEX-4T1 based 5047 bp STh-GST fusion, $Amp^R$ | (4) |

**Table S2. antibodies used in these studies**

| supplemental table 2 antibodies used in these studies | | | | |
|---|---|---|---|---|
| antibody | description | source | RRID* | reference(s) |
| 9A6 | mouse monoclonal IgG1 kappa light chain against human CEACAM6 | Santa Cruz sc-59899 | | (5) |
| A0115 | rabbit polyclonal anti-human CEA (CEACAM5) isolated from hepatic metastasis of colon carcinoma | Dako, Denmark A0115, (now Agilent A0115) | AB_2335697 | |
| 611-1322 | goat IgG (H&L) anti-rabbit IgG (H&L)-HRP conjugated | Rockland 611-1322 | AB_219723 | |
| GST 3-4C | Mouse IgG2b,kappa monoclonal clone GST 3-4C raised against recombinant GST. | Invitrogen 13-6700 | AB_2533028 | |
| GST-STh | Rabbit polyclonal antibody raised against GST-STh fusion protein | Fleckenstein lab | | (4) |
| Anti-O78 | rabbit polyclonal antibody against O78 lipopolysaccharide | Penn State E. coli Reference Center | | (6) |
| A11008 | Goat anti-rabbit IgG (H&L) cross-absorbed, Alexa Fluor 488 conjugated secondary antibody | ThermoFisher | AB_143165 | |
| A-11072 | F(ab')2-Goat anti-Rabbit IgG (H+L) cross-adsorbed, Alexa Fluor 594 conjugated secondary antibody | ThermoFisher | AB_2534116 | |
| 115-205-166 | Goat anti-mouse IgG (H&L) 12nm colloidal gold affinity-purified conjugate | Jackson ImmunoResearch | AB_2338734 | |
| 111-215-144 | goat anti-rabbit IgG (H&L) 18 nm colloidal gold affinity purified conjugate | Jackson ImmunoResearch | AB_2338017 | |
| anti-dmLT | polyclonal sera from mice vaccinated with dmLT | Fleckenstein lab | | (7) |
| Anti-IAP | Polylonal rabbit IgG raised against recombinant (aa3-206) intestinal alkaline phosphatase | Invitrogen PA5-2210 | AB_11153471 | |
| Lysozyme | Polyclonal Rabbit IgG raised against human lysozyme | Invitrogen PA5-16668 | AB_10984852 | |
| CD9 | Mouse monoclonal C-4 vs amino acids 101-210 of human CD9 | Santa Cruz Biotechnology sc-13118 | | |
| Anti-mouse IgG | Affinity-purified horse anti-mouse IgG (H&L) conjugated to HRP. | Cell Signaling Technology 7076 | AB_330924 | |
| * https://antibodyregistry.org | | | | |

### Table S3 ECV-associated proteins with increased abundance in LT-treated enteroids

| supplemental table 3. ECV-associated proteins with increased abundance in LT treated-enteroids. | | | | |
|---|---|---|---|---|
| protein | description | known/putative function(s) | expression | references |
| FCGBP | mucin-like exoprotein with repetitive vWD domains | mucosal defense?; covalently binds to MUC2 via vWD domains | goblet cells | (8, 9) |
| CLCA1 | secreted, non-integral membrane protein | abundant in secreted mucus protein | goblet cells | (8, 10) |
| CD59 | GPI-anchored membrane protein | associated with exosomes; prevents formation of complement membrane attack complex | ECVs | (11, 12) |
| MUC2 | major mucin secreted by goblet cells | mucosal barrier to pathogens including ETEC | goblet cells, mucin | (13-16) |
| PRSS8 | membrane-anchored serine protease | potential role in epithelial barrier function[a] | epithelia including small intestine | (17) |
| CEACAM6 | GPI-anchored membrane glycoprotein protein | intercellular adhesion; binding to E. coli via type 1 fimbriae | intestinal epithelia, granulocytes | (18-20) |
| MUC1 | membrane bound mucin glycoprotein | epithelial surface mucin; glycocalyx component. Potential bacterial decoy | intestinal epithelial surfaces, exosomes | (16, 21, 22) |
| HP | haptoglobin preprotein | potential antibacterial activity | exosomes | (23, 24) |
| BAIAP2L1 | membrane adapter protein | microvillus biogenesis elongation; | intestinal brush border. | (25, 26) |
| TMBIM1 | transmembrane | ? | endosomal membranes | |
| MUC12 | membrane bound mucin glycoprotein | epithelial surface mucin; glycocalyx component. | intestinal epithelial surfaces | (8) |
| S100A6 | cytoplasmic intracellular $Ca^{++}$binding protein | $Ca^{++}$ sensing? | ubiquitous cytoplasmic protein | (27) |
| CTSZ | cathepsin Z | lysosomal cysteine proteinase, involved in killing of Salmonella in macrophages | ubiquitous cytoplasmic protein | (28) |
| EPS8L2 | cytosolic | thought to link growth factor stimulation to actin organization/cytoskeletal remodeling | expression in intestinal epithelial cells[b], exosomes | (23) |
| YWHAE | cytosolic | Adapter protein implicated in the regulation of a large spectrum of both general and specialized signaling pathways. Binds to a large number of partners, usually by recognition of a phosphoserine or phosphothreonine motif. | ubiquitous expression, exosomes | (23) |
| FABP1 | cytosolic protein fatty acid binding | binds long-chain fatty acids/bile acids | enriched in intestine (proximal enterocytes), liver | (29, 30) |
| UGP2 | cytosolic UDP-glucose pyrophosporylase | produces UDP-glucose | distal enterocytes | (31) |
| CKB | creatine phosphokinase | catalyzes the transfer of phosphate between ATP and various phosphogens such as creatine phosphate | intracellular, distal enterocytes | |
| OLFM4 | Olfactomedin 4 | surface glycoprotein involved in multiple signaling pathways | intestinal stem cells, transit-amplifying cells | (32-36) |
| CEACAM5 | Carcinoembryonic antigen (CEA) | intercellular adhesion molecule, mediates homophilic and heterophilic interactions with CEACAM6 | distal enterocytes, Paneth cells, goblet cells[c] | (37) |

[a]https://www.proteinatlas.org/ENSG00000052344-PRSS8
[b] https://www.proteinatlas.org/ENSG00000177106-EPS8L2
[c] https://www.proteinatlas.org/ENSG00000105388-CEACAM5

**Table S4. Original data and figure elements in Figshare repository**

| Supplemental table 4. Datasets and original figures | | |
|---|---|---|
| title | type | DOI (https://doi.org/... |
| Supplemental Dataset 1 | Mass spectrometry data | 10.6084/m9.figshare.22812755 |
| Figure 2D | TEM | 10.6084/m9.figshare.25641060 |
| Figure 2E | TEM | 10.6084/m9.figshare.25641096 |
| Figure 2F | TEM | 10.6084/m9.figshare.25641117 |
| Figure 2G | TEM | 10.6084/m9.figshare.25641183 |
| Figure 3A | TEM | 10.6084/m9.figshare.22814159 |
| Figure 3B | TEM | 10.6084/m9.figshare.22814207 |
| Figure 3C | immunoblot | 10.6084/m9.figshare.24302596 |
| Figure 3E inset IAP | immunoblot | 10.6084/m9.figshare.22806857 |
| Figure 3E inset CEACAM | immunoblot | 10.6084/m9.figshare.22806866 |
| Figure 4A LT ECV dotblot | Immunoblot | 10.6084/m9.figshare.25193030 |
| Figure 4C ECV input IAP | Immunoblot | 10.6084/m9.figshare.25191584 |
| Figure 4C ECV input CEACAM6 | Immunoblot | 10.6084/m9.figshare.25192799 |
| Figure 4C ECV pulldown LT | Avidin_HRP blot | 10.6084/m9.figshare.25193009 |
| Figure 4C toxin +/- controls | Avidin_HRP blot | 10.6084/m9.figshare.25193036 |
| Figure 4G SEC immunoblot | Immunoblot | 10.6084/m9.figshare.25383097 |
| Figure 6A | Immunogold TEM | 10.6084/m9.figshare.26255087 |
| Supplemental figure 2 | Immunogold TEM | 10.6084/m9.figshare.25731456 |
| Supplemental figure 3 | TEM image | 10.6084/m9.figshare.22814150 |
| Supplemental figure 3A | TEM image | 10.6084/m9.figshare.24302620 |
| Supplemental figure 3A | Immunoblots anti-CD9 | 10.6084/m9.figshare.26100613 |
| Supplemental figure 3A | Immunoblots anti-CEACAM6 | 10.6084/m9.figshare.26102722 |
| Supplemental figure 3A | Immunoblots anti-lysozyme | 10.6084/m9.figshare.26102782 |
| Supplemental figure 5C | Anti-CEACAM6 immunoblot | 10.6084/m9.figshare.26356780 |
| Supplemental figure 5C | Anti-OmpF immunoblot | 10.6084/m9.figshare.26356723 |
| Supplemental movie S1 | ppt presentation | 10.6084/m9.figshare.24302626 |

**Table S5 EV protein yields**

| sample | source | amount of starting material | EV protein concentration[a] (mg/ml) |
|---|---|---|---|
| diarrheal stool | human | 7.5 ml[b] | 0.2 |
| fecal pellet | mouse | 346 mg extracted in 15 ml of buffer | 0.3 |
| culture supernatant | small intestinal organoid (Hu235D) | 7.0 ml | 0.1 |
| [a]per ml of SEC fraction determined by Qubit Protein Assay<br>[b]pooled 1.5 ml of sample from each of 5 patients | | | |

**Movie S1. ETEC are eliminated with CEACAMs**
Schematic (left) outlines the challenge of CEABAC10 mice with GFP-expressing ETEC: H10407(pGFPmut3.1), and fractionation of CEACAMs/bacteria from fecal pellets. Movie (right) is compiled from a Z-stack of confocal images of bacterial/CEACAM complexes remaining in the pellet.

**Dataset S1. Tandem mass tag mass spectrometry (TMT-MS) of ECV.** Complete dataset for ECV from untreated and LT-treated enteroids.
